# Epigenetic and transcriptional landscape of heat-stress memory in woodland strawberry (*Fragaria vesca*)

**DOI:** 10.1101/2023.05.26.542514

**Authors:** María-Estefanía López, Béatrice Denoyes, Etienne Bucher

## Abstract

We have previously reported that different stresses can lead to substantial DNA methylation changes in strawberry. Here, we wanted to assess the heritability of heat-stress induced DNA methylation and transcriptional changes following asexual and sexual reproduction in a plant. Woodland strawberry (Fragaria vesca) is an ideal model to study epigenetic inheritance in plants because it presents two modes of reproduction: sexual and asexual. Here we wanted to assess if heat-stress induced DNA methylation changes can be transmitted via asexual reproduction and whether past stresses can also affect sexually propagated progenies. Our genome-wide study provides evidence for a memory acquisition and maintenance in F. vesca. We found that certain DNA methylation changes can stably be transmitted over at least three asexual generations. Finally, the first sexual generation by selfing from stressed maternal and their respective non-stressed asexual daughter plants showed both shared and unique stress responses. This suggests that an acquired molecular memory from the previous heat-stress event was transmitted. This molecular memory might be involved in functional plasticity and stress adaption, an important aspects that will have to be investigated in future studies. Finally, these findings may contribute to novel approaches that may contribute to the breeding of climate-ready plants.

**IN A NUTSHELL:** *Background:* With ongoing climate change, natural plant populations and crops are facing stress situations more frequently and at higher intensity. These unfavorable growing conditions force plants to develop strategies to adapt to persist. One of these strategies involves epigenetic mechanisms which can affect the activity of genes without altering the actual DNA sequence. These molecular modifications can be retained by plants as a molecular “memory” which might be used later to better respond to a stressful event.

*Question:* Is there multi-generational persistence of heat-stress induced epigenetic patterns in strawberry and are heritable epigenetic changes associated with stress adaptation?

*Findings:* We found that the strawberry methylome and transcriptome respond with a high level of flexibility to heat-stress. In addition, we took advantage of the two reproductive modes of strawberry (asexual and sexual) to evaluate the acquisition and maintenance of molecular stress memory. We showed how specific DNA methylation and gene expression changes can persist for a long time in progeny plants. We found that the asexual, and seemingly also sexual progenies can retain information in the genome of a past stressful condition that was encountered by its progenitor.

*Next steps:* Our work presents valuable epigenetic and transcriptional screening data to understand plant memory maintenance and transmission over generations. The most important next step will be to assess if heritable stress-induced epigenetic changes can contribute to stress adaptation through a plant competition experiment in natural environments.

**One sentence summary:** Strawberry can transmit molecular stress-memory at the DNA methylation and transcriptional level over multiple generations which may play an important role in stress adaptation.

## Introduction

Crop production and food security are threated due to global climate change (IPCC, 2022). Therefore, looking for sustainable and efficient agricultural strategies is important for safeguarding worldwide food security. High variations in temperature, water availability, changes in light quality, and nutrients availability result in a hostile growth environment for plants. In order to adapt to such changes, plants depend on life strategies which involve alterations at the genome level (Zhang et al., 2021). For example, floral transition modification is one of the life strategies plants deploy. In contrast to the single flowering event in annual plants, perennials (plants that live more than two years) can flower and restart vegetative growth every year (Battey, 2000; Wang et al., 2009). Many perennial herbaceous plants have two types of reproduction: sexual and asexual (vegetative). These include plants such as cassava (*Manihot esculenta*), greasewood (*Larrea tridentata*), blackbrush (*Coleogyne ramosissima*) and woodland strawberry (*Fragaria vesca*) (Yang and Kim, 2016). Plants that reproduce *via* asexual reproduction produce genetically identical ramets or daughter plants from roots, rhizomes, stems (including stolons, which are elongated stems), tubers, leaves or even inflorescences (Price and Marshall, 1999; García-Verdugo et al., 2013). Compared to sexual reproduction, asexual reproduction results in a reduction of genetic diversity, and thus a loss of adaptative capacity over time (García-Verdugo et al., 2013).

This could be due to these plant’s short-distance dispersal strategy resulting in a limited avoidance of adverse environmental conditions (McKey et al., 2010). However, asexually propagated plants can differ from the parental plant due to the accumulation of random genetic mutations (somaclonal variation) contributing to phenotypic variability (Burian et al., 2016; Jiang et al., 2014). Besides somaclonal genetic variations in an asexual population, epigenetic changes can also perturb the function of the genome and can thus contribute to heritable phenotypic variation (Wibowo et al., 2018). In the case of the perennial *F. vesca*, the extension of lateral meristems form stolons (St) that can produce lateral buds to generate a complete daughter plant (Wang et al., 2019). It was shown that soil grown asexual *F. vesca* daughter plants have low genetic mutation rates (∼ 0.6 mutations per daughter plant), compared to *in vitro* tissue culture, where alterations of DNA methylation levels increase both somaclonal genetic and epigenetic variation frequency (Wang et al., 2019; Cao et al., 2021; Bairu et al., 2011). Previous studies explored DNA methylation levels associated to the climate of origin, such as temperature and altitude, of natural plant populations, including *F. vesca*. (Guarino et al., 2015; Liang et al., 2014; Sammarco et al., 2022; De Kort et al., 2020; Galanti et al., 2022; Dubin et al., 2015). On the other hand, our latest work showed how dynamic the epigenome of *F. vesca* can be at DNA methylation level when plants were exposed to a broad panel of stress conditions (López et al., 2022). The stable inheritance of these epigenetic marks or epimutations might be important contributors to the adaptation and biological diversity of the asexual and sexual progenies. For instance, phosphate starvation and drought stress in rice cultivars generated short-term memory in newly grown leaves that presented altered DNA methylation levels associated to stress response genes (Kou et al., 2021; Secco et al., 2015). Similarly, successive drought exposures in clonal white clover (*Trifolium repens*) induced methylation changes which were maintained across several asexual progenies (Rendina González et al., 2018). Jasmonic (JA) and salicylic acid (SA). Treatments that induced DNA methylation changes in dandelions were also efficiently transmitted (between 74% to 92%) to the following cycle of asexual reproduction (Verhoeven et al., 2010). For Arabidopsis, it has been shown that hyperosmotic stress induced gain and loss of DNA methylation can be heritable to following generations through the maternal line (Wibowo et al., 2016). Here, we focused on heat-stress because climate projections predict temperature increases and extreme heat waves, which will lead in a conflict between water for human and agriculture consumptions (NOAA, 2022). We have previously demonstrated that heat-stress in *F. vesca* was of particular interest from an epigenetic point of view because it resulted in the most drastic genome-wide and local loss of DNA methylation compared to all other tested stresses (López et al., 2022). Here, we wanted to study how heat-stress impacts DNA methylation and transcripts accumulation and how this information can be transmitted to following generations. In addition, we investigated if stress-induced phenotypic changes can be inherited. Overall, we identified DNA methylation changes at cytosines in symmetric sequence contexts (CG and CHG) that can be stably inherited over asexual reproduction cycles. Conversely, we found that even though heat-stress resulted in vast DNA methylation changes at asymmetric cytosines, the original DNA methylation patterns were re-stablished to the original state particularly at the second asexual reproduction cycle. Our comparative epigenomic and transcriptomic analyses suggest that extensive variations in DNA methylation (partially associated to gene expression) and transcription in parental lines during heat-stress is sufficient to generate a “molecular memory” in the sexual and asexual progenies which might contribute to stress adaptation.

## Materials and methods

### Plant growth and material

In this investigation we utilized the same homozygous near-isogenic line (NIL, Fb2:39–47), *F. vesca cv. Reine des Vallées* (RV), showing perpetual flowering and runnering (process to produce stolons) as described and sequenced previously (Urrutia et al., 2015; López et al., 2022). Sterilized seeds were germinated in water on Whatman filter paper for two weeks and transferred to 1/2 Murashige & Skoog (MS) medium (Duschefa cat# M0222), 30% sucrose, and 2% phytagel (Sigma-Aldrich cat# P8169). Plants were then grown for 4 weeks in long-day conditions (16 h light 24°C/8 h dark 21 °C) in growth chambers (Panasonic, phcbi: MLR-352/MLR-352H) prior to stress, as described in our previous study (López et al., 2022).

### Stress assays in vitro

One-month-old seedlings were transferred to a fresh MS media and growth chambers at 24°C/21°C (day/night),16 h light/8 h dark, as control conditions. For heat-stress, plants were exposed to 30°C (day/night) for one week followed by 2 days of recovery (24°C/21°C) on fresh medium as well as the control plants. Then, the plates were transferred to 37°C (day/night) for 1 week followed by 2 recovery days (Figure 1A). We sampled arial parts of plants for the molecular analyses. To reduce variability resulting from individual plants, three biological replicates of 5 pooled plants were collected per condition. Samples were harvested in 1.5 mL tubes between 9:00-11:00 a.m. and immediately frozen in liquid nitrogen and stored at -80°C.

**Figure 1.**
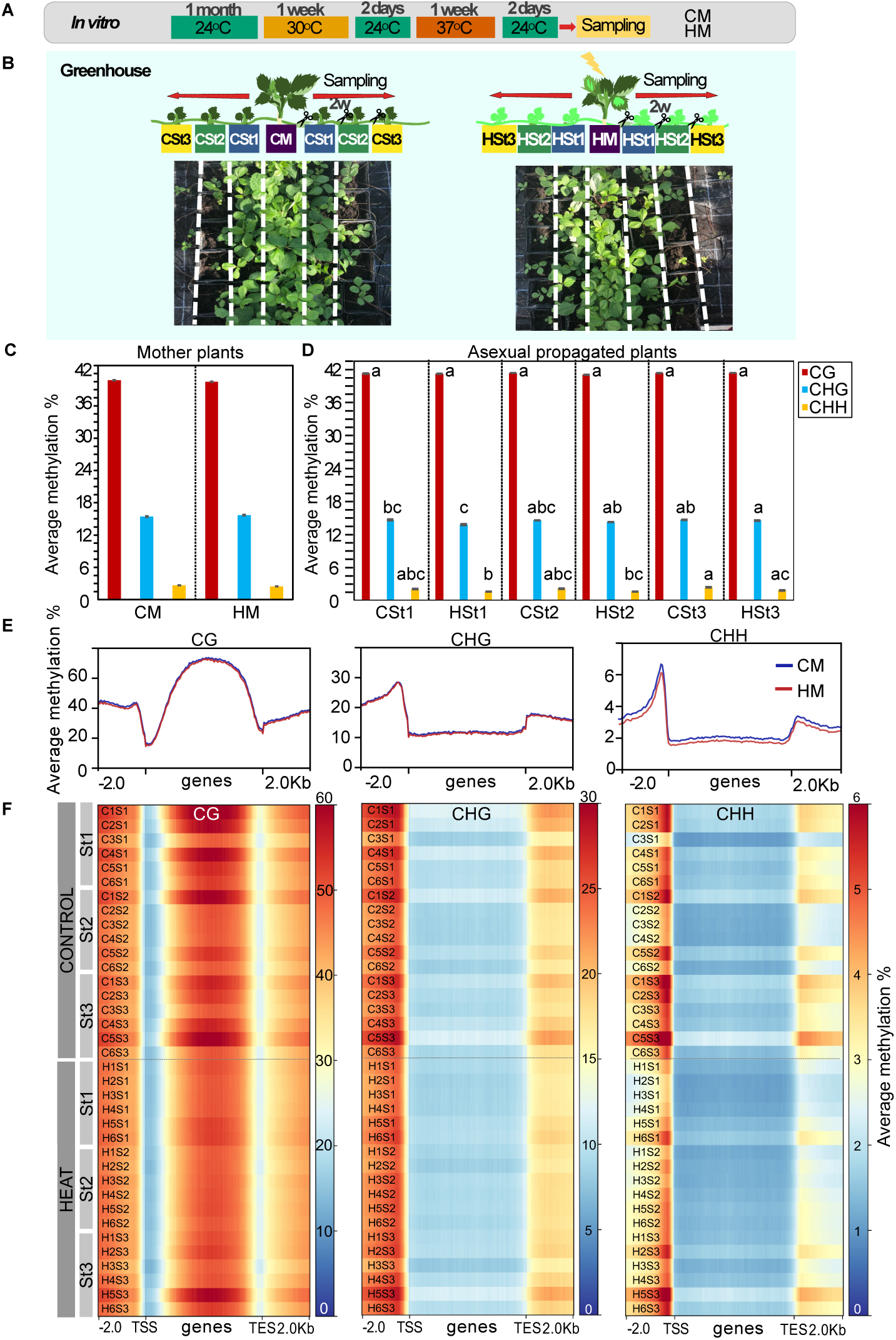
DNA methylation variability after heat-stress and asexual reproduction of F. vesca. **(A)** Scheme of experimental design with treated and untreated plants. One-month-old strawberry seedlings treated with heat-stress *in vitro*. CM: control mother, HM: Heat-stressed mother. **(B)** Asexual reproduction via stolons (St) in the greenhouse. Photographs show 2-weeks-old daughter plant produced by HM and CM plants. St1, St2 and St3: daughter plants of the first, second and third asexual generations respectively: CSt1from CM, CSt2 from CSt1 and CSt3 from CSt2; HSt1 from HM, HSt2 from HSt1 and HSt3 from HSt2. Average of DNA methylation levels for each cytosine context (CG, CHG, CHH) between **(C)** control and heat-stress mother plants (no statistical difference); **(D)** asexual daughter plants per condition (only common cytosines positions among all samples were considered that had a minimum coverage of 5 reads). Means not sharing a common letter at a given asexual group are significantly different (Tukey’s Test; P < 0.05). (E) Plots showing distribution of DNA methylation (top: mCG, middle: mCHG, and bottom: mCHH) around genes of plants treated with (HM) and without stress (CM). **(F)** Heat maps showing the distribution of DNA methylation around genes of all daughter plants from each asexuall generation and their replicates (n=6). Average methylation percentage (within a sliding 50-bp window) was plotted 2 kb upstream of TSS, over the gene body and 2 kb downstream of TES.

### Greenhouse propagation assays

After heat and control treatment the *in vitro* plants were transferred to soil (one plant per pot) in square plastic pots (size: 12x12x10 cm) and to a greenhouse with long day conditions (24°C/21°C day/night and 60%-70% humidity). Twelve mother plants (M) from control (CM; n=12) and heat-stress (HM; n=12) conditions were propagated clonally on three generations (asexual reproduction) (Figure 1B). From each mother plant, the two first stolons (St1) were kept producing one daughter plant each in individual pots. After two weeks, following root formation in St1, the stolons were cut to get independent plants from their mother plant (M). Next, from each daughter plant issued from the first cycle of asexual reproduction (CSt1 and HSt1; respectively daughter plants emerged from M placed in control or in heat stress conditions) one stolon per plant was maintained to get the second cycle of asexual reproduction (St2). Similarly, after two weeks the stolon was cut to get an independent daughter plant; CSt2 and HSt2 from CSt1 and HSt1. We kept the daughter plants CSt2 and HSt2 to generate the third asexual cycle of vegetative reproduction (St3) and after 2 weeks the stolon was cut to maintain independent daughter plants; CSt3 and HSt3 (Figure 1B). All plants were kept until their reproductive phase (∼ 4-month-old) to collect seeds. For molecular analyses, the second new fully extended leaf from each independent daughter plant (2 plants per condition and per cycle) were sampled at an interlap of two weeks for each asexual reproductive cycle. For bisulfite- and RNA-sequencing, we pooled the leaves from the two daughter plants coming from the same mother plant (n=6). The samples were harvested in 1.5 mL tubes between 9:00-11:00 a.m. and frozen in liquid nitrogen. All collected samples were grinded and separated in two tubes to obtain RNA and DNA from the very same samples.

### DNA extraction and whole-genome bisulfite sequencing (WGBS) analysis

Genomic DNA from strawberry plants was extracted with the DNA isolation, plant DNA mini kit, peqGOLD (VWR Life Science, cat# 13-3486-01). For DNA library preparation and bisulfite sequencing, samples were sent to Novogene (Hongkong, China). Paired-end reads were obtained on an Illumina (150 bps) NovaSeq6000 instrument. The DNA methylation datasets were analyzed using the Epidiverse/wgbs pipeline (Nunn et al., 2021b) with the parameters previously reported in López et al., 2022. Lambda DNA reads were used to evaluate the bisulfite conversion rate. An average of 61,257,191 reads (∼ 30X coverage) were produced per sample, of which 81% mapped properly to the *F. vesca* genome of the line that we previously sequenced (López et al., 2022). The average non-conversion rate among the samples was 0.28% (See **Supplemental Table S1** for more details). To calculate global methylation ratios, we use the parameters previously described in (López et al., 2022). Global DNA methylation levels were computed by combing all bedGraph files into a unionbedg file pre-filtered for a minimum coverage of 5 reads per cytosine position and per sample. Only cytosine positions in common in all samples were kept. R-package ggplot2 v.3.3.5 was used for the visualization plots and unpaired Student’s t-test for statistical analysis. p<0.05 was selected as the point of minimal statistical significance in all the analyses.

### Identification of differentially methylated regions (DMRs)

The previously pre-filtered bedGraph files obtained were used as input for the EpiDiverse/dmr bioinformatic analysis pipeline to identify DMRs (Nunn et al., 2021a) with default parameters (**Supplemental Figure S1**). Global DNA methylation and DMR plots were performed with R-package ggplot2 and deepTools v.3.5.0. We produced genome browsers tracks with DMRs in our publicly accessible JBrowse instance: https://jbrowse.agroscope.info/jbrowse/?data=fragaria_sub.

### RNA-seq analysis and definition of differentially expressed genes (DEGs)

Total RNA extractions were performed using NucleoSpin RNA Plus, Mini kit for RNA purification with DNA removal column (cat# 740984.50). *In vitro* mother plant samples (CM; MP; n= 6) and daughter plant samples (St1; St2; St3; n=36) were sent for Illumina paired end read sequencing (150 bp) to Novogene (Hongkong, China). RNA-seq analyses were performed as previously described in (Roquis et al., 2021; López et al., 2022). Briefly FastQC v.0.11.9 and trimmomatic v.0.39 packages were used for quality control and trimming. Salmon v1.4.0 was used as sequence mapper. DESeq2 package was use for the quantitative differential gene transcription analysis at the European Galaxy platform with default parameters (**Supplemental Figure S2**).

**Figure 2.**
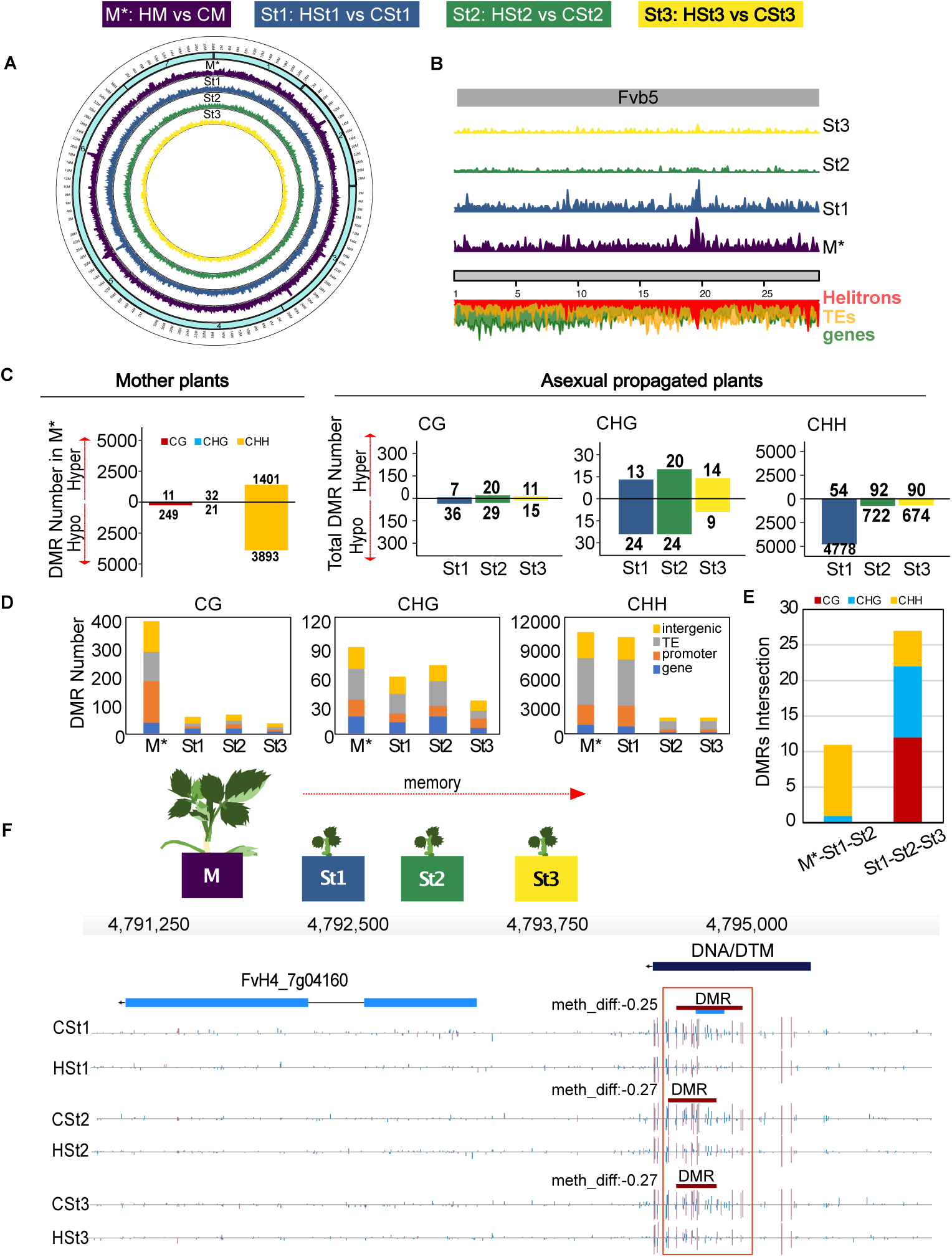
Transmission of heat-stress-induced differentially methylated regions (DMR)s in *F. vesca* during asexual reproduction. **(A)** Circos plot showing Genome-wide distribution of DMR densities on all 7 strawberry chromosomes (turquoise boxes) for each asexual plant generation. M*: Mother ; St1, St2 and St3: daughter plants of the first, second and third asexual generations respectively. M*: HM and CM, St1:HSt1 and CSt1, St2:HSt2 and CSt2, and St3:HSt3 and CSt3 comparisons (*: samples collected *in vitro*). **(B)** DMR density depicted for chromosome 5 (Fvb5) for all asexual generations. Below, the chromosome is indicated in grey, green shows gene density, yellow the TE density and red *Helitron* density along this chromosome. **(C)** Total number of stress-induced hyper-(hyperDMRs) and hypomethylated DMRs (hypoDMRs) are separated by sequence context. Differences between HM and CM *in vitro* are shown in the left plot. Daughter plants comparisons are shown in the right plot. **(D)** Distribution of DMRs in genomic features: promoter, gene body, transposable elements (TE) and intergenic regions. Minimum overlap required: 1bp. **(E)** Bar plot depicting counts of common DMR locations (minimum overlap: 1bp) containing hypo- and hyperDMRs per context among all populations. Boxes above the plot indicate the color codes for DNA methylation: red for CG, blue for CHG and yellow for CHH sequence contexts. **(F)** Genome browser views of overlapped DMRs located in promoter region of PR5-like receptor kinase gene (FvH4_7g04160) in St1, St2, St3. Depicted are genes structures (top panels, UTRs in light blue, exons in blue), TEs (dark blue) and DNA methylation levels (histograms). Boxes above the histograms indicate identified DMRs with methylation difference ratios (color code for DNA methylation: red for CG).

Genes showing a DMR (promoter and gene body located) and DEGs were annotated using Gene Ontology (GO) and Kyoto Encyclopedia of Genes and Genomes (KEGG) annotation downloaded from the Genome Database for Rosaceae (GDR) (Jung et al., 2019). Genes containing hypo- and hypermethylated DMRs were classified to investigate potential functions through GO enrichment by AgriGOv1.2 (Tian et al., 2017). Similarly, all DEGs were extracted with p-adj≤0.05 for the GO functional analysis with Fisher’s exact test p-values and Hochberg (FDR) as multi-test adjust method (≤ 0.05 as the cut off). To identify potential enriched pathways, KEGG analysis was performed with the R-package: clusterProfiler (Wu et al., 2021); pvalueCutoff = 0.05; pAdjustMethod = "fdr". Plots were generated with the R-package ggplot2.

### Phenotyping assays

Five-month-old asexual daughter plants of the three asexual generations from control and heat-stress mother plants were treated with an ascending gradient of temperature (30°C, 35°C, 40°C) to evaluate possible phenotypes and adaptation traits. Plants from the greenhouse were moved to plant growth chambers (Panasonic, phcbi: MLR-352/MLR-352H) for heat-stress treatment (n=10 per condition). Maximum quantum yield of primary photosystem II photochemistry (QY_max_= *F_V_*/*F_M_*) was measured at each temperature with a chlorophyll fluorescent device (FluorPen FP 110, Photon Systems Instruments PSI). Maximum PSII quantum yield was measured from dark-adapted leaves. *F_M_* is the maximum fluorescence intensity emitted by dark-acclimated samples *F_V_* is measure by the variable fluorescence increment that is due the transition from dark-adapted state with all-open reaction centers to the all-closed state during saturating flash of light (FluoroPen manual and user guide). At the same time, leaf chlorophyll content was measured with a chlorophyll meter, SPAD-502Plus (KONICA MINOTA), for 3 trifoliate opened leaves form each plant. Selected leaves were marked with a string to follow the variation of the chlorophyll content over the increase of temperature.

A new group of younger daughter plants (one-month-old, n=12 per condition) from a replicate experiment were exposed to 37°C and 42°C for 24h to evaluate flowering time.

### Transcriptional memory assays

To evaluate the behavior of first generation (S1) obtained after sexual reproduction by selfing, seeds form mother and daughter plants were collected. Seedlings were grown as previously described under stress assays *in vitro*. One-month-old seedlings from each group (CM, HM, CSt1, HSt1) were exposed to 37°C for 24h and then moved to recovery conditions (24°C) for two days. For transcriptome analysis pools of five seedlings were sampled with three biological replicates from each group at time 0 without stress and at 6h stress and after 2 recovery days.

## Results

### Global stress-induced DNA methylation differences across clonally propagated strawberry plants

In our previous study, we described how a large set of abiotic stresses, including heat-stress, resulted in changes in DNA methylation patterns in treated *F. vesca* plants (López et al., 2022). We found that heat-stress had the strongest impact on the epigenome, leading to a genome-wide loss of DNA methylation. To assess whether heat stress-induced DNA methylation changes can be maintained after asexual reproduction, we compared the methylation profiles of heat-stressed (H) and non-heat-stressed (control, C) mother plants (M) and the profiles of their daughter plants produced by asexual reproduction and grown under standard conditions (**Figures 1A, 1B**). Mother plants were heat stressed at the seedling stage as detailed in (López et al., 2022). In summary, two cycles of stress were applied for one week each (30°C and then 37°C) on *in vitro* seedlings (**Figure 1A**) and a part sampled. Remaining seedlings were planted in greenhouse. Asexual reproduction was then performed via stolons (St) to get daughter plants. Three successive asexual generations were obtained via stolons (St) to get daughter plants named St1, St2 and St3. Eventually, we obtained eight types of plants: control mother plant (CM, *in vitro*), heat-stressed mother plant (HM, *in vitro*), CM-derived daughter plants: CSt1, CSt2, CSt3, and HM-derived daughter plants: HSt1, HSt2, HSt3. All daughter plants were grown in the greenhouse. To assess genome-wide DNA methylation levels and transcriptional changes at all generations at the same developmental stage, DNA and RNA from the second new trifoliate leaf from each daughter plant was extracted. First, DNA samples were submitted to whole genome bisulfite sequencing (WGBS, 30x genome coverage) (**Supplemental Table S1**). We carried out a global quantification of DNA methylation (m) in the three sequence contexts: CG (mCG), CHG (mCHG) and CHH (mCHH). We used the same parameters as previously described in López *et al*., 2022 to filter out the cytosine positions without sequencing coverage and to calculate average DNA methylation levels. In HM *in vitro*, the global DNA methylation levels after heat-stress decreased by 1% and 8% for the CG and CHH contexts compared to CM (no statistical difference). Conversely, global DNA methylation increased by 2% in the CHG context (**Figure 1C**). Even though HSt1, HSt2, and HSt3 showed some variability in global DNA methylation compared to the controls, these global differences were not statistically significant (**Figure 1D**). To assess methylation variation in genic and non-genic sequences, we screened the methylome data in all three sequence contexts separately in three regions: 2 kb upstream of genes, along the gene body, and 2 kb downstream of genes using 50-bp sliding windows (**Figures 1E, 1F**). We observed loss of mCHH in gene bodies, transcription start (TSS) and end (TES) sites in HM vs. CM. Meanwhile mCG and mCHG were quite stable in both HM and CM groups (**Figure 1E**). Furthermore, methylation levels showed an overall higher variability between CM-derived daughter plants compared to the HM-derived daughter plants (**Figure 1F**). We selected genes with body methylation (gbM) as described by (Bewick et al., 2019). Genes containing gbM after stress (HM) showed low mCG and mCHG variability (**Supplemental Figure S3A**). HSt1, HSt2, and HSt3 presented hypermethylation in gene bodies in the CG context. For the CHH context, hypomethylation observed at TSS and TES in HM plants, was moderately maintained in asexual progenies (**Supplemental Figure S3A**). To document the variation in the profiles of DNA methylation over transposable element (TE) body and flanking regions, we compared the methylomes from HM vs. CM, and between their daughters (St1, St2, St3) on the *F. vesca* TE annotation. We found that different TE families exhibited distinct DNA methylation dynamism in TE body and flanking regions after heat-stress especially at asymmetric cytosines (**Supplemental Figure S3B**).

### Heat-stress causes long-term changes in genomic DNA methylation patterns

To explore the dynamics of DNA methylation at specific loci in detail, we assessed differentially methylated regions (DMRs) for each sequence context using the Epidiverse Toolkit with default parameters (Nunn et al., 2021a). Samples clustered according to each treatment group separating the control progenies from the heat-stress progenies (**Supplemental Figure S1**). We further compared the methylomes of the HM daughter plants with the methylomes from CM daughter plants from the same generation: M* (HM vs. CM) (*: samples collected *in vitro)*, St1 (HSt1 vs. CSt1), St2 (HSt2 vs. CSt2) and St3 (HSt3 vs. CSt3) in the greenhouse. We observed that the genome-wide distribution of DMR densities was enriched at specific genomic regions preferentially near centromeric and pericentromeric regions after heat-stress over the seven chromosomes in the *in vitro* parental line comparison “M*” (**Figure 2A**) as described in López et al., 2022. The heat stress treatment performed on the mother plant (M) allowed to conserved DMR hotspots on the three asexual generations not submitted to this stress (**Figure 2B**). This effect was stronger on St1 and decreased with the asexual generation level. In general, the distribution of DMRs did not seem to follow a clear gene or TE distribution pattern in the genome; except for its correlation with *Helitron* transposable element distribution (**Figure 2B**) (López et al., 2022). Overall, we observed a steady reduction of DNA methylation differences over each asexual reproductive cycle.

To classify the DMRs, we counted the statistically significant DMRs and identified whether they were hypomethylated regions (hypoDMRs) or hypermethylated regions (hyperDMRs) for each of the three sequence contexts (**Figure 2C**). The majority of identified DMRs were detected in the *in vitro* mother plants after heat-stress with 5607 DMRs. Reduced numbers of DMRs were detected in the following asexual progenies, 4912 in St1, 907 in St2, and 813 in St3 (**Figure 2C**). Most DMRs were hypoDMRs (98% in St1; 85% in St2; 86% in St3) relative to the control condition in all sequence contexts and in all groups of plants. To test whether genic and/or non-genic regions were enriched in DMRs, we qualified DMRs based on their intersection with the following annotations: promoters (empirically defined as 2 kb to 50 bp upstream of a TSS), gene bodies, and intergenic regions (minimum overlap required 1bp). Many of the CG-DMRs in M* (37%) were in promoters, while in the St progenies most of the CG-DMRs (between 29-36%) were in gene body and intergenic regions. The highest proportion of CHG- and CHH-DMRs (between 35%-50%) were in TEs in M* and daughter plants (**Figure 2D**). In order to identify potential functional roles of the DMRs located in genic regions (genes and promoters), we performed a singular enrichment analysis (SEA) using the AgriGOv2 tool (Tian et al., 2017) (see Materials and Methods). HM vs. CM seedlings showed 3187 DMRs over genic regions. The 2745 genes with a DMR (over body and promoter) were enriched in transport activity, establishment of localization and in cell membrane (**Supplemental Figure S4A**). We did not find any significant enrichment for DMRs associated with genes in the St groups vs. M*. To further investigate the role of heat stress in inducing stable epigenetic changes in the three asexual generations, we analyzed whether the DMRs present in the progenies were acquired directly from the heat-stressed parents. Even though each daughter plant is genetically identical, they differed at the epigenetic level showing unique and common DMRs between St1, St2, St3 and M* (**Figure 2E**). Looking at all generations together (M*-St1-St2-St3) no DMRs were observed that were directly induced in M* by heat-stress and maintained over the three clonal generations. However, the analyse of each asexual generation with the heat stress mother plant M* highlighted the first asexual generation maintained the highest number of DMRs (267) compare to the two following asexual generations St2 and St3. These DMRs were mostly in CHH context (**Supplemental Figure S4B**). In addition, 11 DMRs overlapped in between M*, St1 and St2 (**Figure 2E**). Because different growing conditions of plants, *in vitro* for mother plants and *ex vitro* for the three asexual generations can affect DMRs, we compared them among the three St generations to identify stable and inherited methylation marks. We found that 47% CG-DMR, 37% CHG-DMRs and 9% CHH-DMRs in St1 were maintained in at least one of the two next asexual generations (**Figure 2E**). Taken together, these results showed that heat-stress can induce specific stable DNA methylation changes that can persist over at least three asexual generations, albeit with a steady loss of DNA methylation changes (particularly in the CHH context) over asexual generation. Notably, 27 DMRs were shared among all St generations (Example in **Figure 2F**).

### Heat-stress induces a stable transcriptional memory maintained over asexual generations

Because stress response impacts the regulation of gene expression, we assessed the relationship between induced changes in DNA methylation and gene expression. First, we identified differentially expressed genes (DEGs) from the exact same samples as those used for BiSeq analysis (**Figures 1A, 1B**). Hierarchical clustering of significant DEGs in at least one group showed that St1, St2, and St3 cluster together apart from the M* group (**Supplemental Figure S4C**). DEGs from M* plants showed higher fold change values compared to the St groups (**Supplemental Table S2**). After heat-stress, *in vitro F. vesca* seedlings (HMvs.CM) showed in total 10935 DEGs of which 57% were down-regulated (**Figure 3A**). The asexual progenies comparisons (HSt vs. CSt) showed: 1156 DEGs in St1, 664 in St2 and 23 in St3. In contrast to the M* plants, the first two progenies had mainly up-regulated DEGs: 85% in St1 and 62% in St2 (**Figure 3A**, **Supplemental Table S3, Table S4**). To explore the role of the heat stress on maintaining the transcriptional response over the three asexual generations, we examined whether the DEGs identified after heat-stress were transcriptionally preserved among the asexual progenies. Comparable with the DMRs, St1 had more DEGs from the stress-exposure of the parents. Notably, 28% up- and 56% down-regulated genes in St1 were in common with those observed in M*. Moreover, 13% up- and 33% down-regulated genes from St2 were shared with those found in M* (**Figure 3A**). St3 presented the least transcriptional changes compared with the first two asexual progenies with no DEGs shared with M* (**Supplemental Table S5**). Even though most transcriptional changes are lost over asexual generations, 15 DEGs were seemingly maintained over two asexual reproductive cycles from M* to St1 and St2 (**Figure 3A**). To better understand the potential roles these DEGs could have in each plant group, we performed gene ontology (GO) enrichment analyses (**Supplemental Figure S4D**). The M* group was broadly enriched in biological process such us metabolic process, response to stimulus, reproduction, macromolecule transport and modification. In addition, it showed enrichment in molecular functions such as transcription factor activity and regulation, and transferase activity (**Supplemental Figure S4D**). Notably, response to stress and stimulus was still present in the DEGs in St1 (**Supplemental Figure S4E**) and metabolic process in St2 (**Supplemental Figure S4F**). Most likely because of the reduced number of DEGs in St3, no significant GO enrichment could be detected. To complement the analysis, we performed a pathway functional analysis (KEGG) from each group. For M* we found enzymes related with reductase, monooxygenase, and transferase activity. St1 included transcription factors, peroxidase, and laccase activity (**Supplemental Figure S4G**). Laccase activity was also detected as a significant pathway in the St2 KEEG analysis. Overall, the data indicated that 4% of the transcriptional heat-stress response can be preserved at the transcriptional level in an immediate asexual generation, here St1 (60 days after the initial heat-stress).

**Figure 3.**
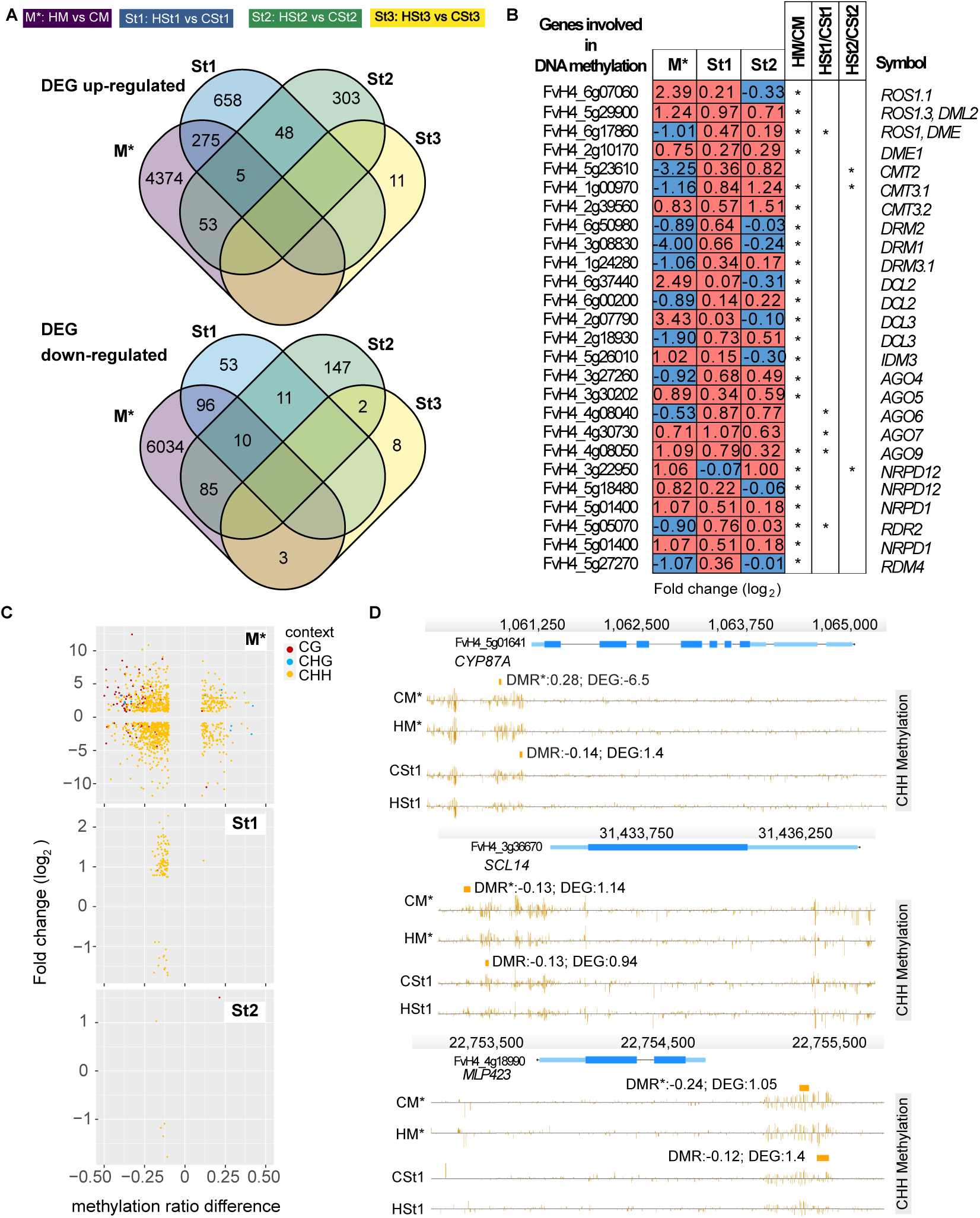
Heat-stress induced differential gene expression in mother plants and in the non-stressed asexual progenies. **(A)** Venn diagrams displaying the intersections of differentially expressed genes of the four groups: after two days of recovery from heat-stress (HM vs CM) *in vitro* and 2-week-old daughter plants (St1, St2, St3) (results from DESeq2, p-adj:<0.05). The top set represents up-regulated genes and the bottom one the down-regulated genes. **(B)** Transcriptional values (fold change in log2) of genes involved in DNA methylation and RdDM pathway. * p-adj:<0.05. First column shows gene ID number and last column shows gene symbols. **(C)** Scatterplots showing DEGs related with DMRs (promoter and gene body) in M* (upper plot); St1 (middle); and St2 (bottom). Points show the relationship between transcript levels (fold change in y-axis) and DNA methylation (methylation difference in x-axis). The color codes for DNA methylation are: red for CG, blue for CHG and yellow for CHH contexts. **(D)** Genome browser views depicting the DEGs after heat-stress in M* and St1. The examples shown here are: cytochrome P450 87A3-like (FvH4_5g01641), MLP-like protein 423 (FvH4_4g18990), Purple acid phosphatase 22 (FvH4_6g18411). Depicted are genes structures (top panels, UTRs in light blue, exons in blue), TEs (red and dark blue) and DNA methylation levels (histograms). Rectangles above the histograms indicate identified DMRs with methylation difference ratios (color codes for DNA methylation: red for CG, blue for CHG and yellow for CHH contexts). Transcription values in fold change (log2; results from DESeq2, p-adj:<0.05). *: samples collected *in vitro*.

### Spontaneous and stable DNA methylation changes are associated with gene expression patterns

In order to investigate possible direct effects of heat-stress on the DNA methylation machinery, we analyzed the expression of genes involved in different DNA methylation mechanisms based in the last genome annotation v4.0.a2 (Li et al., 2019). Genes coding for proteins involved in DNA demethylation (e.g., ROS1 orthologues) were differentially expressed. Certain DNA methyltransferases (CMT2, CMT3) were downregulated in M*; however, they were up regulated in St1 and St2 (**Figure 3B**). In addition, members of the *ARGONAUTE* gene family were also differentially expressed. *AGO4* and *AGO6* were down regulated in M* but in St1 *AGO6*, *AGO7* and *AGO9* were up regulated. The non-catalytic subunit common to nuclear DNA-dependent RNA polymerases II, IV and V (NRPD12) was up-regulated in M*. *RNA-dependent RNA polymerase 2* (*RDR2*) and DICER-like genes were mostly up-regulated only in M* (**Figure 3B)**. The observed changes in transcript levels of genes involved in several DNA methylation processes suggests an ongoing adjustment of DNA methylation over successive asexual reproductive cycles.

To investigate the relationship between transcription and DNA methylation, we compared plants from the same generation: M* (HM vs. CM) (*: samples collected *in vitro*), St1 (HSt1 vs. CSt1), St2 (HSt2 vs. CSt2) and St3 (HSt3 vs. CSt3) in the greenhouse. We identified 492 DEGs (5% of the total DEGs) in M* that presented a DMR in their promoter or gene body regions, 94 DEGs in St1 (8%), and 16 DEGs in St2 (2%). We did not detect a DEG in St3 that was associated to a DMR. The presence of hypo- or hyperDMRs in CHH context did not correlate with the observed gene expression patterns (up- or down-regulation, **Figure 3C**). For instance, CG hypoDMRs were mostly related with up regulated genes. For instance, 89 DEGs in M* were associated to a CG-DMR. Of these 65 up-regulated DEGs correlated with hypoDMRs in promoter (**Supplemental Table S6**). Only 9 DEGs were related to CHG DMRs in M* (**Figure 3C, Supplemental Table S6**). However, CHH hypoDMRs were equally distributed between up- and down-regulated genes in M, St1 (**Supplemental Table S7**) and St2 (**Supplemental Table S8**).

In addition, to evaluate gene expression with specific hyper- or hypomethylation profiles, we took DEGs with DMRs maintained from M* to St1. We identified 11 genes which kept the DNA methylation difference and transcriptional profiles in M* and St1. Figure 3D shows candidate genes which might be epigenetically regulated and that could be involved in stress memory. Cytochrome P450 family 87 gene (FvH4_5g01641, *CYP87*) had a CHH-DMR in the promoter region after heat-stress in M* with a methylation ration of 0.28 and fold change of -6.5 (log_2_). Conversely, in St1 plants *CYP87* was associated with a hypoDMRs in the CHH context and an increased transcript level (**Figure 3D**). On the other hand, because differences in DNA methylation persisted over more that 60 days after heat-stress (**Figure 2E**), we wanted to assess the extent of potential transmission of changes in DNA methylation and transcription through asexual reproduction in the independent daughter plants. To evaluate whether the 27 stable DMRs in the asexual progenies were related to transcriptional changes, we compared the DMR locations in the vicinity of genes. We found 18 genes related with one of the 27 DMRs; however, these genes did not present significant expression differences. Conversely, 10 genes related with the 27 inherited DMRs were differentially transcribed in M*. To illustrate, in **supplemental Figure S5**, we show three examples where the heat-stress treatment resulted in the up- and down-regulation of genes in M*, and during asexual reproduction differences in DNA methylation were acquired and maintained. These results suggest that the transcription of stress response genes may lead to long-term methylation fluctuations which can be established as a long-term memory.

### Testing transgenerational inheritance of heat-stress induced phenotypic traits in *F. vesca*

To evaluate the extent to which heat-stress could induce transmittable phenotypic changes, we followed the development of HM and CM until adult stages in greenhouse conditions (**as previously described** in López et al., 2022, **Figure 1A**). Following recovery, HM plants showed an early flowering phenotype compared to CM plants (**Figure 4A**). HM plants showed early flowering phenotype and produced a significant higher number of flowers (16 flowers) compared to CM plants (9 flowers) (**Figures 4A, 4B**). However, HSt1, HSt2 and HSt3 asexual generations did not differ in flowering time as compared to control asexual progenies. These results suggest that the observed phenotypical impact of heat-stress is transient and not transmitted over asexual reproduction without stress.

**Figure 4.**
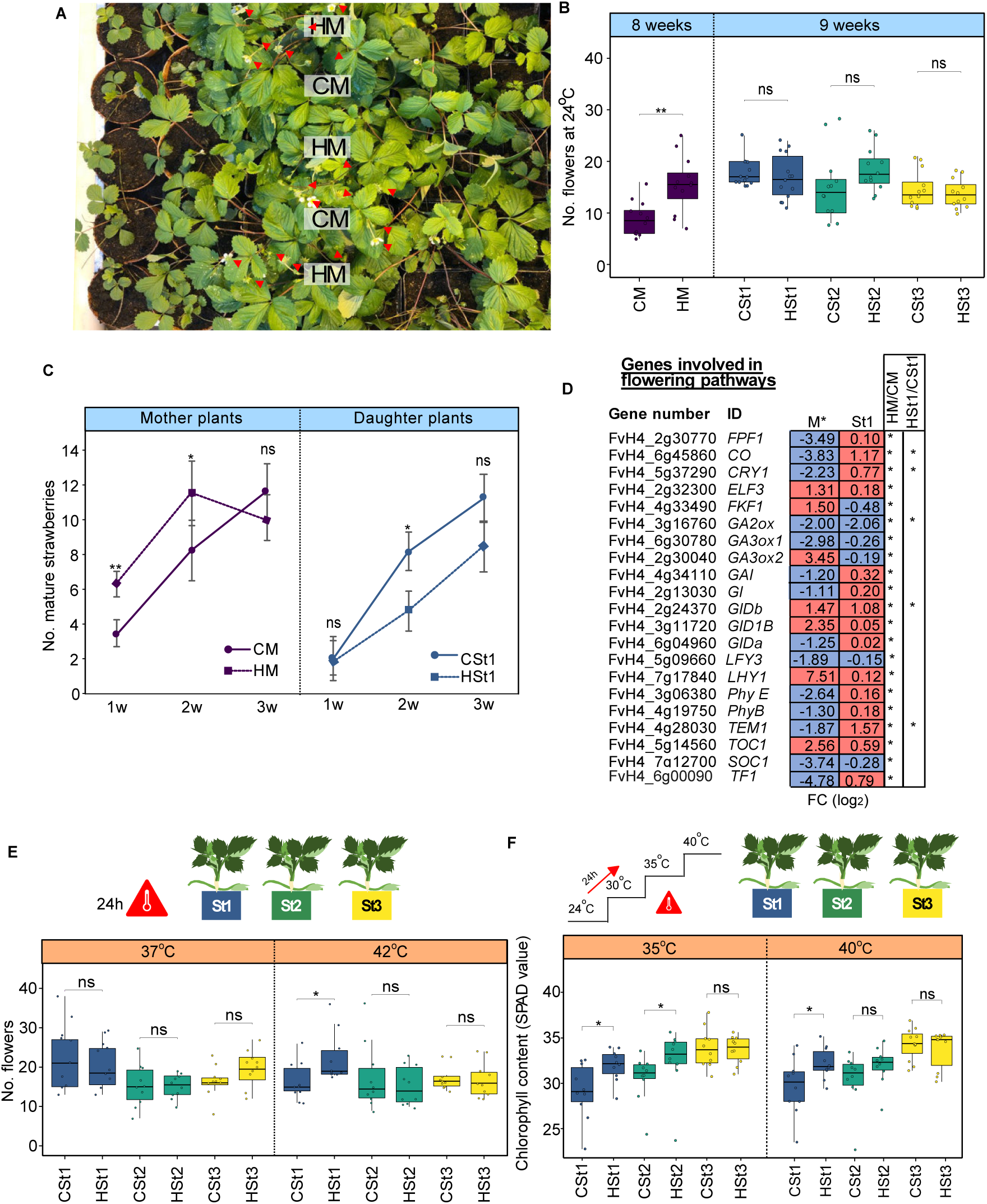
Phenotypes assessed over three clonal generations of plants issued from heat-stressed mother plants. **(A)** Representative photograph of four-month-old mother plants from control (CM) and heat-stressed (HM) in greenhouse during flowering time. Red arrows point closed and opened flowers. **(B)** Boxplots of total numbers of flowers per plant until the first ripe fruit was harvested in the greenhouse. **(C)** Number of harvested strawberries during three consecutive weeks in M and St generations. **(D)** Table showing transcript levels (fold change in log2) of genes linked to flowering including photoperiod and gibberellin pathways in M* (samples collected *in vitro*) and St1 (* adjusted P-value < 0.05, as determined using the DESeq2 **(E)** Total number of flowers per daughter plant exposed to a 24h heat treatment (37°C and 42°C). **(F)** Total leaf chlorophyll content by the measurement of the Soil Pant Analysis Development (SPAD) values by chlorophyll meter of plants submitted to an ascending temperature gradient (+5°C every 24h). All statistical analyses were performed with the Wilcoxon rank sum tests: *P ≤ 0.05; ns: no significant. HM: heat-stressed mother plant; CM: control mother plant; St1, St2 and St3: daughter plants of the first, second and third asexual generations respectively.

**Figure 5.**
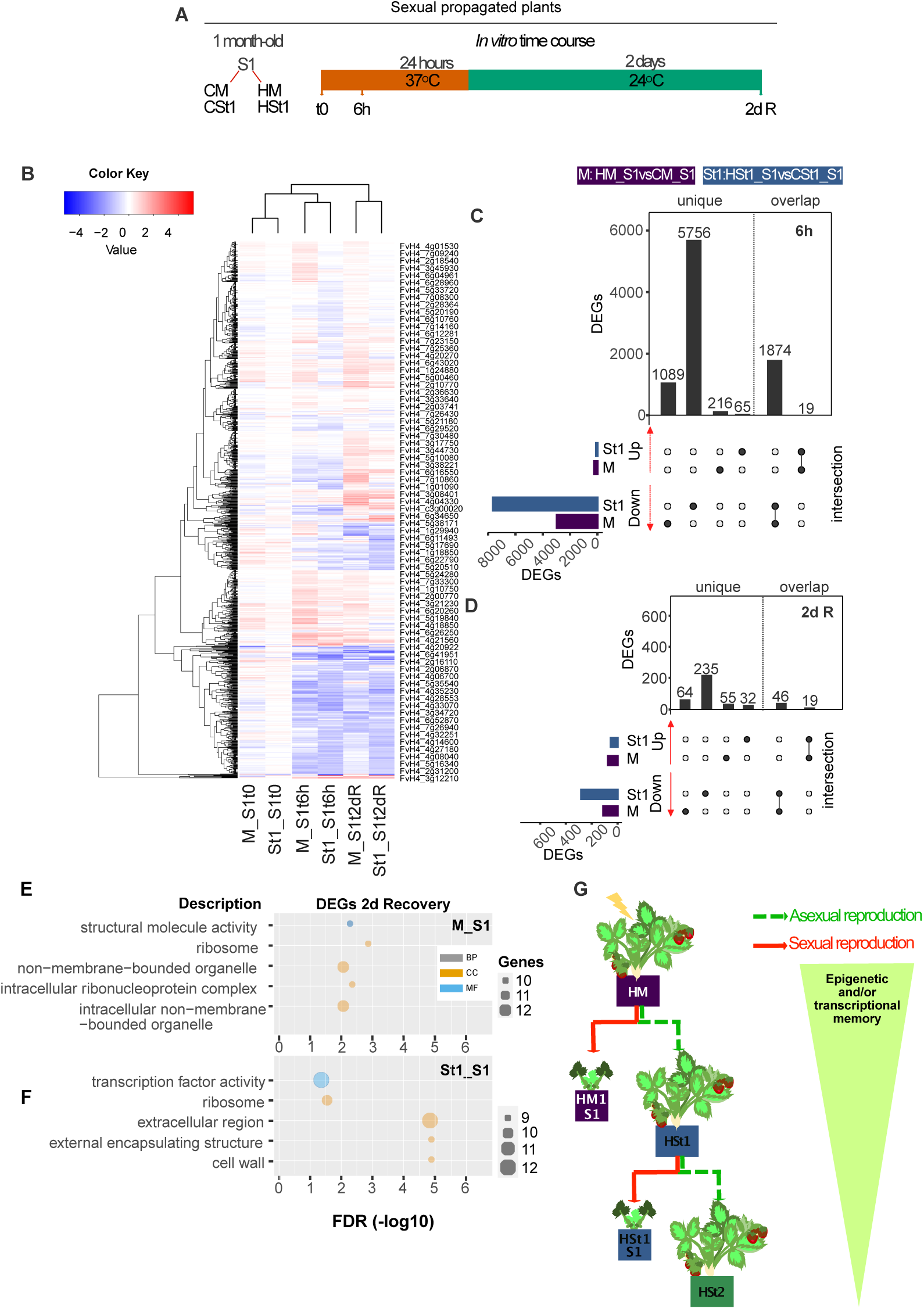
Maternal effects from heat-stress transmitted during sexual reproduction (S1). **(A)** Time course scheme of heat stress treatment on S1. **(B)** Heatmap with DEGs identified in at least one comparison: the M and St groups: M_S1 (HM_S1 vs. CM_S1) and St1_S1 (HSt1_S1vs. CSt1_S1), at time point 0 (t0); after 6 hours (6h) at 37°C; 2 days after recovery (2dR) at 24°C. The upper clustering shows the link among samples from different first generations based on DEGs. Total number of DEGs: **(C)** plot at 6h 37°C exposure, **(D)** plot after two days of recovery (2dR) from heat-stress (results from DESeq2, p-adj:<0.05). Black bar plots show the total number of DEG shared among conditions. Singular enrichment analysis (SEA) of DEGs in M_S1 **(E)** and St1_S1 **(F)** after 2 days of recovery from heat stress. For the bubble plots in **(E,G)** the x-axis indicates the p-adj values. The y-axis indicates the most enriched GO terms in three categories: biological processes (BP, grey), cellular component (CC, orange), and molecular function (MF, blue). The bubble sizes indicate the number of genes in each category. **(G)** Epigenetic and transcriptional memory model through vegetative propagation and sexual reproduction.

As another phenotypic trait we counted the number of mature strawberries to identify alterations in fruit production. Differences appeared at the first two weeks of observation in M* while no difference was observed in St1. We harvested significantly more ripe strawberries from HM during the first and second week after flowering; however, by the third week the number of harvested fruits was reduced and comparable to CM (**Figure 4C**). During the second week CSt1 presented a higher number of fruits compared to HSt1. For the next two asexual cycles of propagation (St2, St3) there were no significant differences (**Supplemental Figure S6A**). Overall, the production of strawberries was similar in all the groups; nonetheless, there was earlier ripening of strawberries in HM which was directly related with the early flowering. In addition, we registered fruit size (width and height) and observed smaller fruits in HM compared to CM (**Supplemental Figure S6B**). Dry biomass and number of seeds (**Supplemental Figure S6C**) were not significantly different among groups. To get more insights into the flowering control in HM and CM, we explored changes in the expression of genes involved in the regulation of flowering (**Figure 4D**). Figure 4D showed only genes that were significantly expressed in at least one group. To illustrate, we found that *CONSTANS* (*CO*) and *TEMPRANILLO* (*TEM1*) were differentially expressed in HM compared to CM *in vitro* conditions and were also differentially expressed in the St1 (HSt1 vs. CSt1) generation. To illustrate, *CO* and *TEM1* were down-regulated in M* (HM vs. CM) and up-regulated in St1 (**Figure 4D**). In addition, we identified genes linked to photoperiod and gibberellin pathways: *CRYPTOCHROME 1* (*CRY1*), *GIBBERELLIN 2-OXIDASE* (*GA2ox*), and *GIBBERELLIN INSENSITIVE DWARF* (*GIDb*) which were significantly differentially expressed in M* and St1. For example, *CRY1* presented a negative fold change in M* while in the St1 generation it showed a significant increase in transcript level (**Figure 4D**). *GA2ox* displayed a significant reduction of expression in HM* and HSt1 compared to controls, meanwhile *GIDb* was highly expressed in both HM* and HSt1. These gibberellin-linked genes conserved expression profiles in M* and St1 suggesting that genes linked to gibberellin pathways may contribute to a short transcriptional memory.

To identify the heritability of heat-stress effects to daughter plants at early stages, we performed a new heat-stress experiment using one-month-old St populations to evaluate differences in flowering time at 37°C and 42°C for 24h (**Figure 4E, see Material and Methods**). We identified early flowering only after 42°C in HSt1 with a total number of 23 flowers compared to CSt1 with an average of 16 flowers per plant (**Figure 4E**). There were no significant differences in St2 and St3 groups. To evaluate the heritability of heat-stress effects to daughter plants at older stages, we submitted St1, St2, and St3 (derived either from HM or CM plants) to a rising gradient of temperature every 24h from 24°C to 40°C (see Material and Methods for details). To assess leaf photochemical efficiency and leaf greenness, we measured chlorophyll content as SPAD (Soil Plant Analysis Development) values (Xiong et al., 2015) (see Material and Methods). We detected significantly higher SPAD values in HSt1 and HSt2 at 35°C and 40°C compared to their corresponding controls (**Figure 4F; Supplemental Figure S6D**). To illustrate, for CSt1 leaves had 29 SPAD at 35°C and 40°C. However, HSt1 leaves had 32 SPAD in average at 35°C and 40°C. In addition, to identify signs of thermotolerance, we evaluated the efficiency of photosystem II (PSII) (Janni et al., 2020; Hassan et al., 2021; Medina et al., 2021). We measured as ratio the maximum quantum efficiency of PSII photochemistry (QY_max_= Fv/Fm), using a chlorophyll fluorometer (**Supplemental Figure S6E**). HSt1 plants showed significantly lower ratios of QYmax at 24°C and 35°C, 0.83 and 0.78 respectively, as compared to CSt1 plants under stress, with 0.85 and 0.82 (**Supplemental Figure S6E**). Nevertheless, we noticed a lower variability at 30°C and 40°C heat treatment in all groups (**Supplementary Figure S6E**). To summarize, only plants of the first asexual generation (HSt1) from heat-treated mother plants (HM) showed significant phenotypic variability compared control plants.

### First sexual generation plant (S1) exhibit transcriptional changes resulting from the maternal heat-stress event

To evaluate a possible adaptive stress response transmitted from a heat-stressed mother plant (HM), we investigated heat-stress response of the transcriptome of the first sexual generation (Selfing 1, S1) from the M and St1 groups: M_S1 (HM_S1 vs. CM_S1), St1_S1 (HSt1_S1vs. CSt1_S1). We performed a short time course where one-month old seedlings were stressed at 37°C for 24h. Expression data was collected at 0h (t_0_), 6h, and 2 days after recovery (**Figure 5A, Supplemental Figure S7A, S7B)**. M_S1 and St1_S1 groups were clustered together in t_0_, 6h and after 2d recovery (**Figure 5B**). At t_0_ there were 30 DEGs in the M_S1 comparison (**Supplemental Table S9**), and 11 in St1_S1 (**Supplemental Table S10**). These results highlight the fact that without external stimuli there are only minor transcriptional differences between the sexual progeny from the control family and the heat-stressed life-history family (**Supplemental Figure S7C**). After 6h of heat-stress, M_S1 displayed 3198 DEGs (**Supplemental Table S11**) and in St1_S1 7714 DEGs (**Figure 5C, Supplemental Table S12**). After 2 days in recovery conditions (24°C) all groups presented less DEGs: 184 DEGs in M_S1 (**Supplemental Table S13**) and 332 in St1_S1 (**Figure 5D, Supplemental Table S14**). To find similar molecular or biological functions among groups, we performed a GO enrichment analysis of the DEGs detected after recovery. DEGs in the M_S1 group were enriched in genes related with cellular components such as ribosome and non-membrane organelles (**Figure 5E**). St1_S1 showed enrichment in genes related to transcription factor activity, cell wall and ribosome (**Figure 5F**). Additionally, we identified 65 DEGs that were in common in M_S1 and St1_S1 after 2 days of recovery from heat-stress (**Figure 5D).** These results suggest that the asexual progenies of HM plants were able to transmit molecular information resulting from the external stimuli that was initially applied to M and being able to be transmitted to sexual reproduction (**Figure 5G**).

## Discussion

Mutations can lead the emergence of favorable and unfavorable traits which by natural selection or breeding will be selected for or against. However, it is now clear that not only genetic mutations but also epimutations can contribute to plant diversity (Quadrana and Colot, 2016; Zamir, 2001; Schmitz et al., 2011). Epigenomes of plants have been examined after unfavorable environmental conditions to evaluate how epigenetic changes might trigger adaptive responses (Verhoeven et al., 2016). High temperature is one of the most studied environmental stress conditions due its negative effects in natural plant populations and crops. Even though epigenetic mechanisms are conserved among plant species (Niederhuth et al., 2016), plants can differ in the way they perceive and respond to high temperatures. For instance, soybean roots and rapeseed seedling under heat treatment showed hypomethylation in all cytosine contexts (Hossain et al., 2017; Gao et al., 2014). We have previously shown, using an extensive panel of stresses, that heat-stress caused the most significant loss of DNA methylation in *F. vesca* which leaded to the reduction of DNA methylation notably in CG and CHH context (López et al., 2022). We confirmed here these observations (**Figure 1C**). In addition, we detected hypomethylation particularly in regions close to TSSs and TESs sites (**Figures 1D, 1E**). Similar results were found in tobacco, maize, and seagrasses where reduction of DNA methylation was mostly identified over promoter regions and gene bodies during heat-stress (Entrambasaguas et al., 2021; Centomani et al., 2015; Qian et al., 2019). Considering the possible positive or negative effects of DNA methylation changes (Seymour and Becker, 2017) and how these epigenetic states are established and inherited are critical aspects to understand stress tolerance and potential stress adaptation (Zion et al., 2020; Nguyen and Gutzat, 2022). We analyzed methylomes from daughter plants issued from three successive asexual generations (St1, St2 and St3) raised in a common unstressed environment. We observed the presence of mainly hypoDMRs in all cytosine contexts particularly in TSS and TES genomic regions (**Figure 2C**). Heat-stress DMRs and DEGs in M* (HMvs.CM) were mostly observed in the first asexual generation, in St1 (HSt1vs.CSt1), with 267 DMRs (**Supplemental Figure S4B**) and 386 DEGs (**Figure 3A**). This first asexual generation is represented by daughter plants produced by the two first stolons of the mother plant, issued from axillary meristems (AXM) (Costes et al., 2014). In the near-isogenic line we used, the two first stolons emerged from the AXM of the 6th adult trifoliate leaf (Tenreira et al., 2017). These AXM were likely not already preformed in the seedlings which displayed three young leaves and in their terminal bud, one leaf in emergence and two or three primordia. Thus, the most likely hypothesis is that the meristematic cells which will become an AXM have kept the memory of the heat stress. This memory fades with each successive asexual generation, which includes a large number of cell division. Studies of patterns of meristematic cell divisions have previously shown how epigenetic and genetic variations occur in cells and how they will contribute to new tissues or a complete plants (Burian et al., 2016; Yao et al., 2021). It has been suggested that mutations that occurred in meristematic cells from the apical meristem will be maintained in the new shoot meristem, whereas changes in subapical meristematic cells are easily displaced and thus finally only affect a small number of cells due to low number of cell divisions (Burian et al., 2016). Similarly, epigenetic variations in meristematic cell can lead to epigenetic mosaicisms during growth and branch development of a wild lavender (*Lavandula latifolia*) which might persit in a ecological and evolutionarily maner (Herrera et al., 2021). Despite the fact that wild type DNA methylation patterns were largely restored, we identified 27 fixed DMRs that were present in all three asexual generations suggesting a transgenerational inheritance of heat-stress induced DNA methylation changes that mostly occurred in the symmetric DNA methylation context (**Figure 2E**). In a similar manner, twelve clonal generations of duckweed (*Lemma minor* L.) exposed to 30°C showed long-term epigenetic memory in CG and CHG loci containing hypermethylated regions (Antro et al., 2022). All these results in symmetric cytosine contexts were comparable with the results obtained in Arabidopsis plants derived from epigenetic mutant lines which displayed stable inheritance of DNA methylation changes in CG and CHG locations compared to CHH (Reinders et al., 2009; Kenchanmane Raju et al., 2019; Jiang et al., 2014). In an induced hypomethylated *F. vesca* population (5-azacytidine treated) it was observed that DNA methylation changes in CG and CHG contexts could be maintained over three sexual reproductive cycles albeit with a notable loss of mCHH patterns (Xu et al., 2016). Furthermore, we identified genes involved in several DNA methylation pathways which were differentially expressed in M*, St1 and St2 (**Figure 3A**) due to heat stress, suggesting that they may contribute to the maintenance of homeostasis of the epigenomes over the generations. It is known that heat-stress triggers rapid responses (from seconds to minutes) and rapid systemic signaling pathways to regulate gene expression (Kollist et al., 2019; Nievola et al., 2017).

In addition, it has been shown in tomato that specific conformation of chromatine states after heat stress leads to over representation of interations between the heat shock transcription factor, HSFA1a, and promoter-enhancers controlling the expression of stress-response genes (Huang et al., 2023). The question remains whether heat stress triggers first transcriptional changes followed by DNA methylation changes, or whether it is the other way around. Phosphate starvation appeared to first activate stress response genes and as consequence DNA methylation differences were induced (Secco et al., 2015). We identified that changes in transcription of genes comparing HM vs. CM plants was concomitant with the formation of stable DMRs in the next three asexual reproductive cycles (**Supplemental Figure S5**); nonetheless, a higher number of DEGs was not linked to DNA methylation changes. Another group of DEGs was associated to changes in DNA methylation in promoters and TESs in genes in M* and maintained until St1 (**Figure 3D**). Altogether, our findings demonstrate that heat-stress has implications in the reduction of DNA methylation levels and gene expression which can enhance the accumulation of molecular signaling to potentially enable a rapid stress response.

Heat-stress has previously been described as a factor which stimulates early flowering, and at the same time, this phenotypic trait can be inherited over several Arabidopsis generations (Balasubramanian et al., 2006; Suter and Widmer, 2013). Stress-induced flowering is considered a survival sign since seed production will guarantee the species prevalence (Takeno, 2016). We confirmed that *F. vesca* submitted to heat-stress had shorter flowering times compared to control plants (Figure 4). A comparable phenotype was observed in the stressed HSt1 generation at 42°C regardless of whether the CSt1 generation was submitted to the heat treatment or not (Figure 4E). Loss of chlorophyll and performance of the photosystem II in *Suaeda salsa*, maize and rice was a phenotype associated with heat tolerance (Kumar et al., 2012; Lu et al., 2003). Damage in photosystem II (PSII) and chlorophyl content due to heat-stress is known to affect productivity of strawberry cultivars (Kadir et al., 2006; Choi et al., 2016). We found that HSt1 at heat-stress maintained the chlorophyl content as compared to CSt1 (**Figure 4F**), suggesting that HSt1 may cope better with subsequent heat-stress.

Despite the maintenance of DMRs and DEGs over asexual reproduction in *F. vesca*, we also wanted to consider inheritance through sexual reproduction. We found different gene expression patterns between progenies (S1) with a heat-stress life event and controls which may be caused by stable DNA methylation modifications (**Figure 2E**). Similar results were found in Arabidopsis in a spaceflight where methylation and transcriptional memory were partially retained over two generations (Xu et al., 2021). In crop plants stress-induced methylation changes were found to often be erased after sexual reproduction (Ganguly et al., 2017; Secco et al., 2015; Zheng et al., 2013). Indeed, epigenetic reprogramming occurs during male and female gametogenesis (Kawashima and Berger, 2014). However, inherited stress-induced epigenetic changes have been observed to primarily be transmitted through the female germline and to be maintained during homologous recombination (Wibowo et al., 2016; Molinier et al., 2006). Common DEGs in *F. vesca* S1 (M_S1 and St1_S1) after heat-stress (**Figure 5**) might be categorized as marker genes to identify lines that went through a stressful life-history. To summarize, our results suggest that heat-stress leads to extensive DNA methylation variability in the mother plant which stays high in the first asexual generation. This variability is then strongly attenuated in the next two asexual generations. However, a proportion of heat-stress induced DMRs appear to be non-random and can be stably inherited over at least three asexual generations of *F. vesca*. It now remains to be tested what defines these non-random regions and if these epigenetic changes can contribute to stress adaptation.

## Conclusions

Sequencing analyses have increased our understanding of genome structure and genome dynamics. And yet, decoding transmission mechanisms of epigenetic characteristics and their dynamism under unfavorable stress conditions remains challenging. Our data provides evidence for a molecular framework of memory acquisition and maintenance in *F. vesca*. Here, we have demonstrated that stressful experiences very early in a plant life can influence the epigenome and transcriptome, and thereby contribute to information transmission. This may have adaptive impacts on both sexual and asexual progenies allowing them eventually to respond more appropriately to subsequent stress conditions. In addition, we described that epigenetic and transcriptomic changes induced by heat stress can be accumulated over time generating a type of molecular memory which seems to have a role in subsequent stress responses. We propose that the early flowering phenotype can be inherited in the first *F. vesca* asexual generation as a “maternal-primed” effect. Altogether, our results suggest that the dynamism of the epigenome and transcriptome in maternal lines are sufficient to alter the signaling and behavior of both asexual and sexual progeny under future stress conditions.

## Data availability

The datasets generated and/or analyzed in this study are available in the Zenodo repository DOI: 10.5281/zenodo.7898322. All the sequencing data from this study have been submitted to European Nucleotide Archive (ENA, www.ebi.ac.uk/ena/, ERP136614). The data can be accessed under the project PRJEB51950. The Bisulfite-sequencing raw read fastq accessions under ERS11389778-ERS11389819. The raw reads of the RNA-sequencing under the accessions: ERS15458182-ERS15458217.

## Acknowledgements

The European Training Network “EpiDiverse” received funding from the EU Horizon 2020 program under Marie Skłodowska-Curie grant agreement No 764965; The European Research Council (ERC) under the European Union’s Horizon 2020 research and innovation program; No 725701 “BUNGEE” to E.B.. Funding for open access charge: Agroscope institutional funding. This study was supported by Agroscope, Nyon-Switzerland. We would like to thank all the members of the EpiDiverse consortium (www.epidiverse.eu) and the Crop Genome Dynamics research group for invaluable support, Eric Remolif for taking care of the plants at the greenhouse.

## Author Contribution

ME.L and E.B conceived the study. ME.L performed the experiments, analyzed sequencing data and wrote the manuscript. B.D provided plant material, designed experiments, and wrote the manuscript. E.B. designed experiments, analyzed data, set up the genome browser and wrote the manuscript.

## Competing interests

The authors declare they have no conflicts of interest.

---

**Figure S1.**
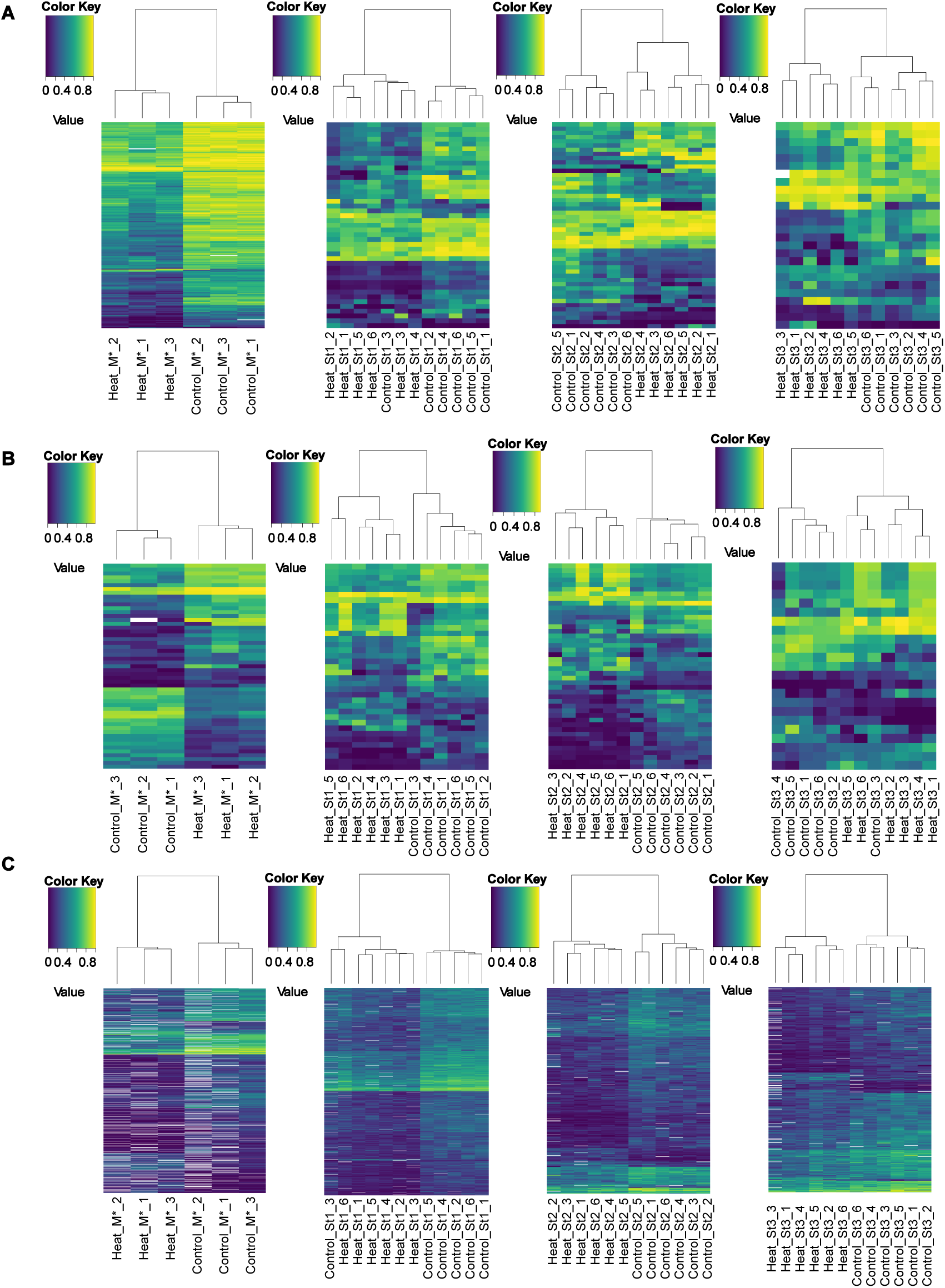
Heatmaps clustering groups based on significant DMRs (q < 0.05) in: (A) CG, (B) CHG and (B) CHH. Methylome comparisons from control mother plants (CM*) against stress mother plant (HM*); control daughter plant (CSt1) vs. heat daughter plant (HSt1) from the first clonal propagation; control daughter plant (CSt2) vs. heat daughter plant (HSt2) from the second clonal propagation; control daughter plant (CSt3) vs. heat daughter plant (HSt3) from the third clonal propagation. *: samples were collected *in vitro*.

**Figure S2.**
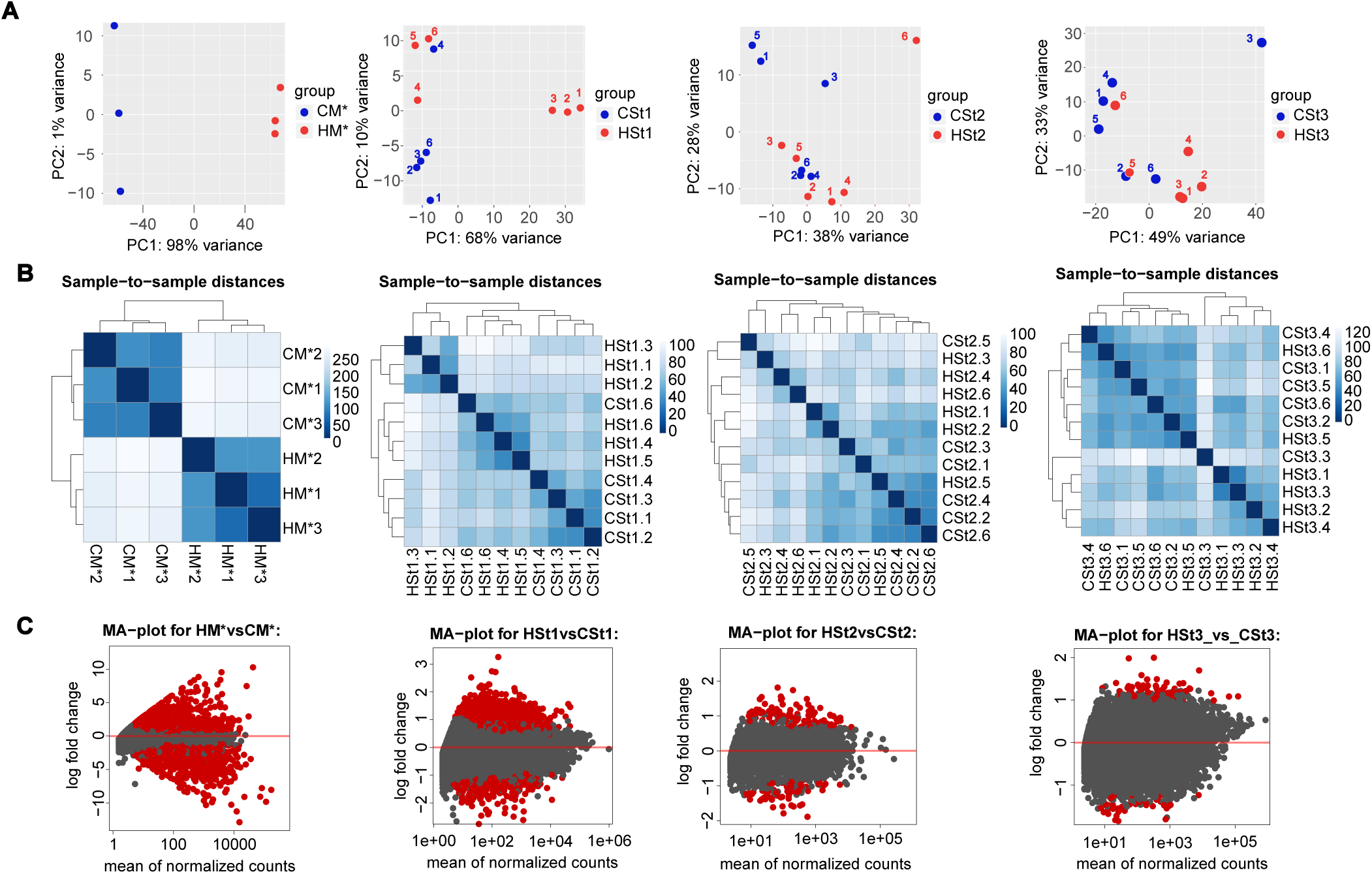
RNA-seq analysis to detect differentially expressed genes (DEGs). (A) PCA plot showing the variability among control and heat-stress samples. (B) Heatmap and cluster showing an overview of similarities and differences among samples. (C) Volcano plot showing the log2 fold change given to the variable mean of normalized counts. Red points indicates if the adjusted p-value is less that 0.1. Plots were obtained by DESeq2 in the Galaxy platform. *: samples collected *in vitro*.

**Figure S3.**
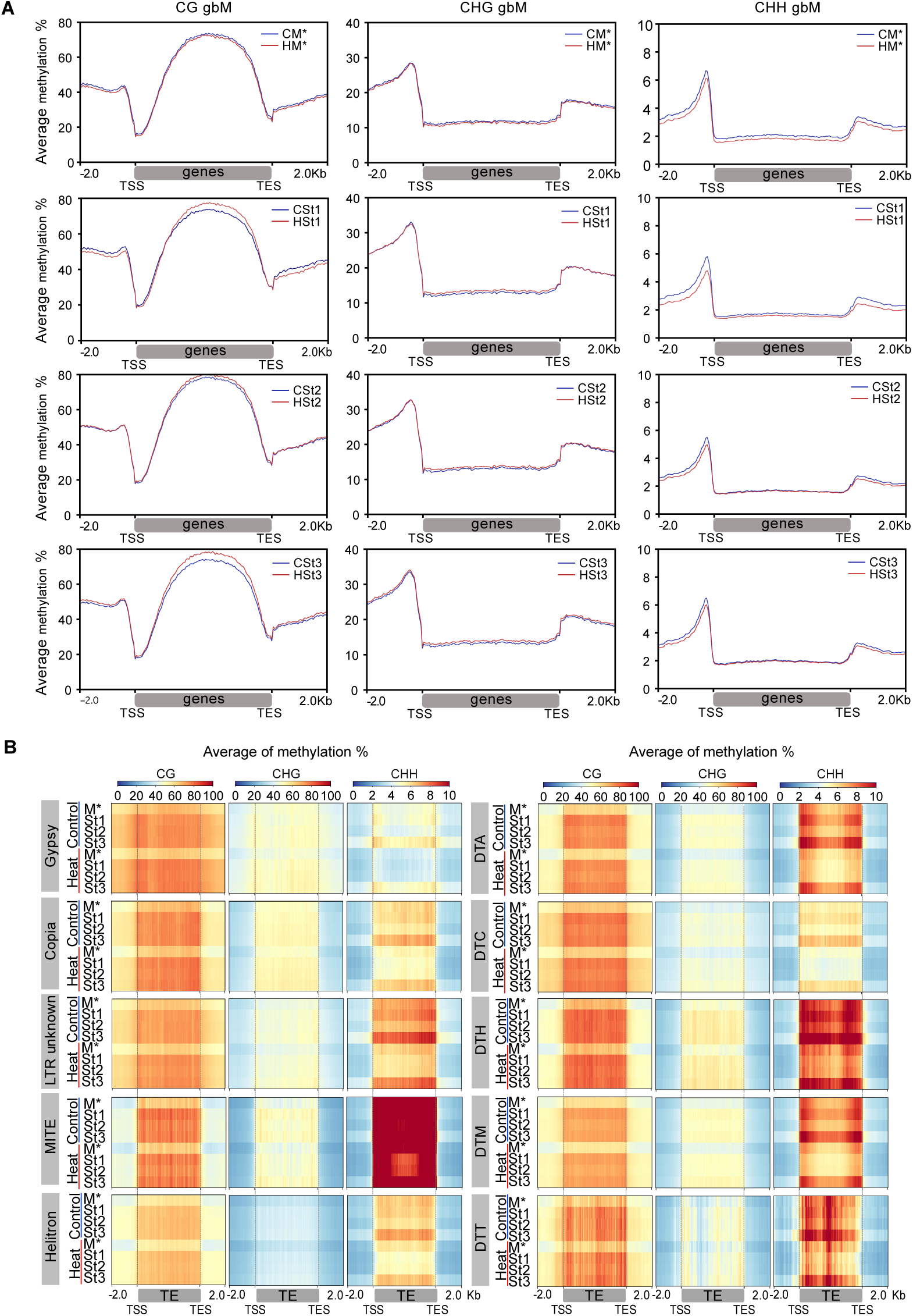
Plot of global DNA methylation profiles at genes with body methylation (gbM). Plots show distribution of DNA methylation in CG (left), CHG (middle) and CHH (right) contexts around genes classified as genes with body methylation (gbM) with and without stress (Control). Mean of the average methylation percentage (within a sliding 50-bp window) was plotted 2 kb upstream of TSS, over the gene body and 2 kb downstream of TES. (b) Heatmaps showing DNA methylation profiles for all the TE families separated by sequence context mCG (left), mCHG (center) and mCHH (right). The mean of the average DNA methylation percentage (within 50 bp sliding windows) was plotted for the TE bodies and 2 kb around the TSS and TES regions. Class I elements (retrotransposons): LTR-Copia, LTR-Gypsy. Class II elements (DNA transposons): TIR: Tc1-Mariner (DTT), hAT (DTA), Mutator (DTM), PIF-Harbinger (DTH), CACTA (DTC); Helitron; Miniature Inverted-Repeat Transposons (MITEs). M*: mother plant (samples collected *in vitro*); St1: Daughter plant of M; St2: Daughter plant from St1; St3: Daughter plant from St2. C: control, H: heat-stress.

**Figure S4.**
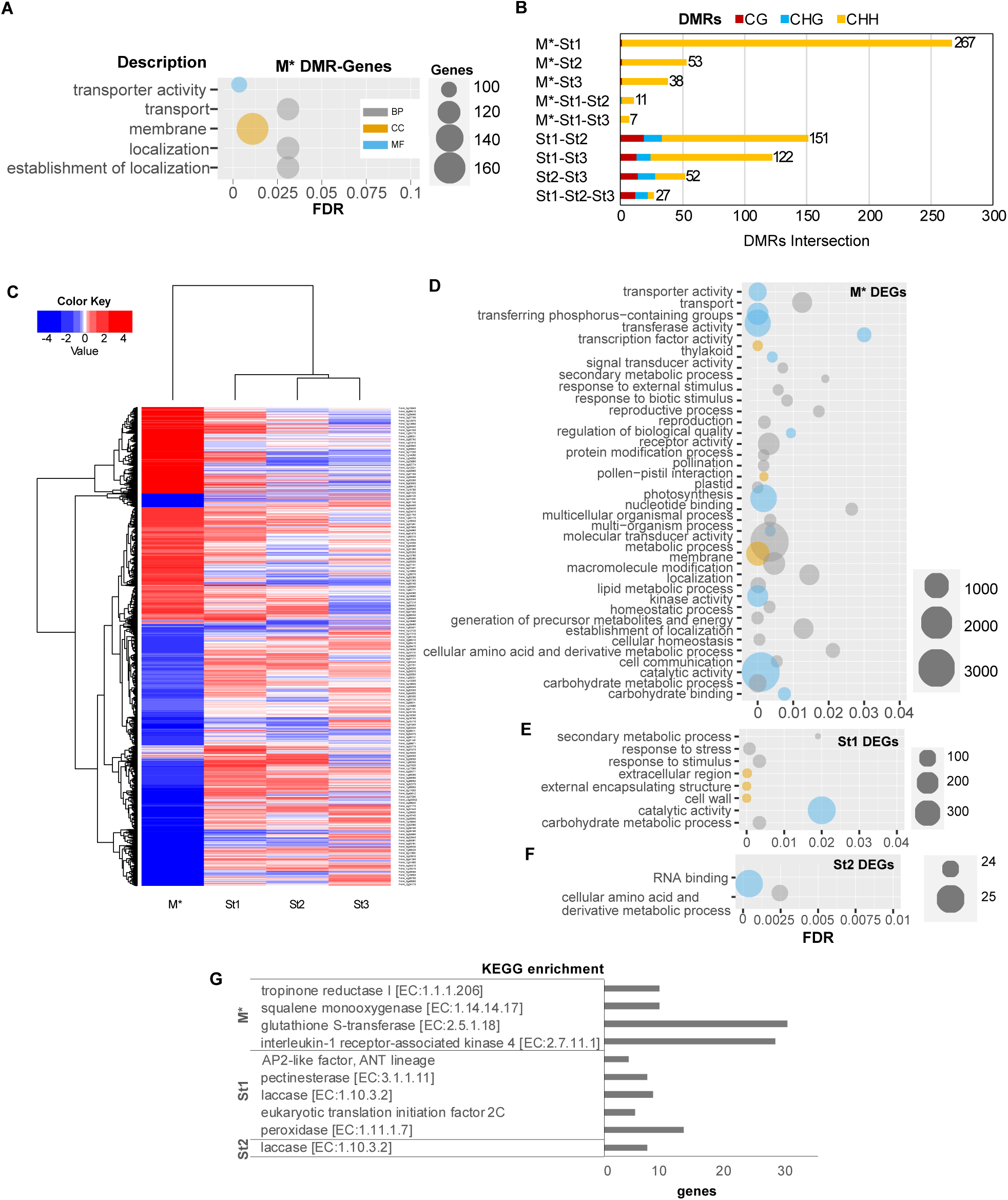
Functional analysis of genes related with a DMR and differentially expressed genes (DEG) in clonal and sexual progenies. Singular enrichment analysis (SEA) of: (A) the total number of genes related with a DMR (in promoter and gene body); (B) Bar plot depicting counts of common DMR locations (minimum overlap: 1bp) containing hypo- and hyperDMRs per context among all populations. Boxes above the plot indicate the color codes for DNA methylation: red for CG, blue for CHG and yellow for CHH sequence contexts. (C) Heatmap with DEGs present in at least one population (M*, St1, St2, St3). Upper cluster shows the link among samples from different clonal generations based on DEGs. (D) DEGs after heat-stress in M*; € DEGs in St1; (F) DEGs in St2, was performed using AgriGOv2 (p-adj:<0.05). (G) KEGG enrichment analysis of the total number of DEGs from M*, St1 and St2 (results from clusterProfiler, p-adj:<0.05). *: Samples collected *in vitro*.

**Figure S5.**
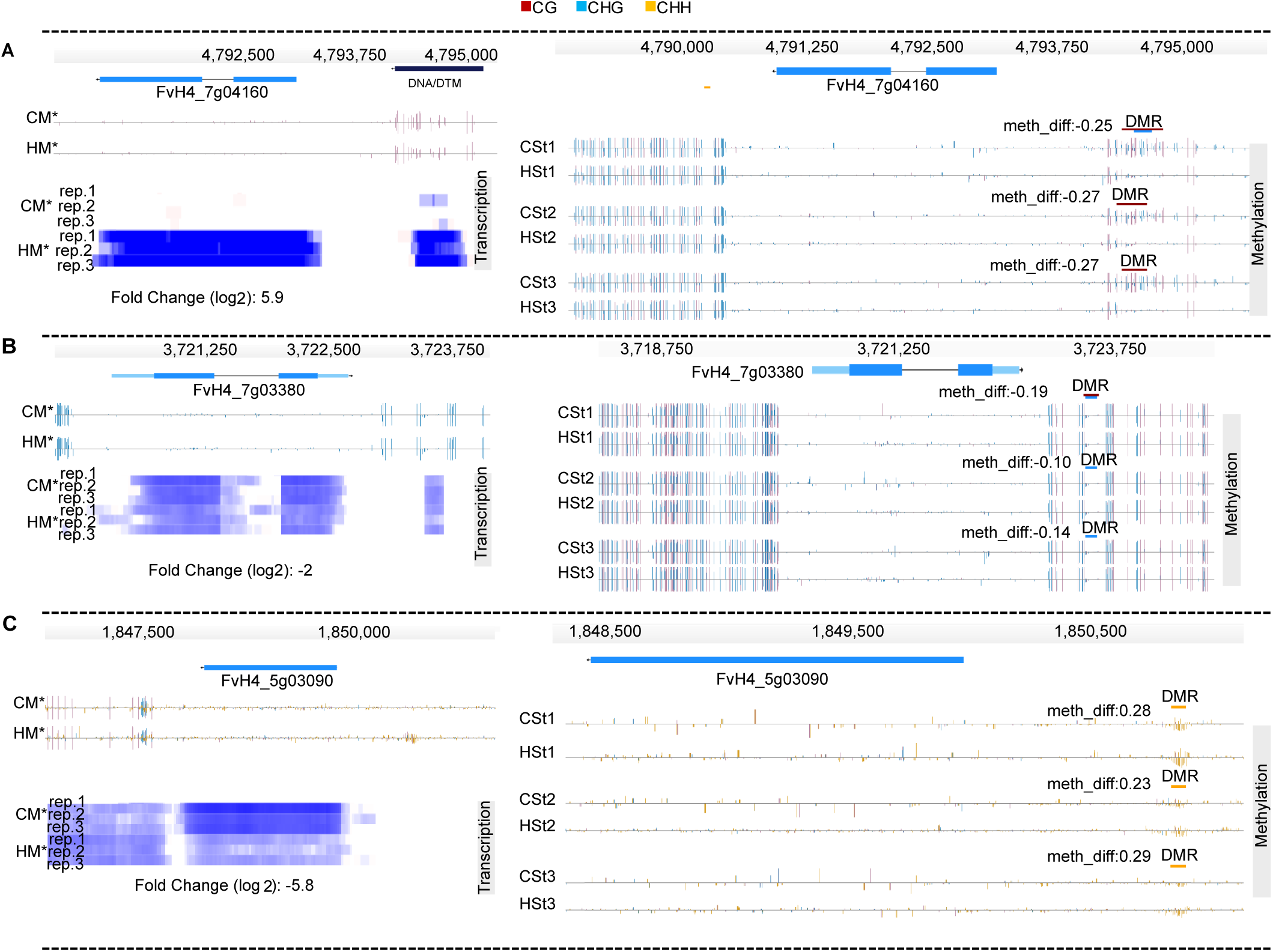
Heat-stress DMRs maintained over three asexual generations (St1, St2, St3) of genes that were differentially expressed in the mother plant (M*). Genome browser views of differentially expressed genes after heat-stress in M* (left) and the DMRs located in promoter regions in St1, St2, St3 at those same genes in the absence of the stress (right). (a) PR5-like receptor kinase gene (FvH4_7g04160); (b) sequence-specific DNA binding transcription factor gene (FvH4_7g03380); (c) uncharacterized gene (FvH4_5g03090). Depicted are genes structures (top panels, UTRs in light blue, exons in blue), TEs (red and dark blue) and DNA methylation levels (histograms). Boxes above the histograms indicate identified DMRs with methylation difference ratios (color codes for DNA methylation: red for CG, blue for CHG and yellow for CHH contexts). *: Samples collected *in vitro*.

**Figure S6.**
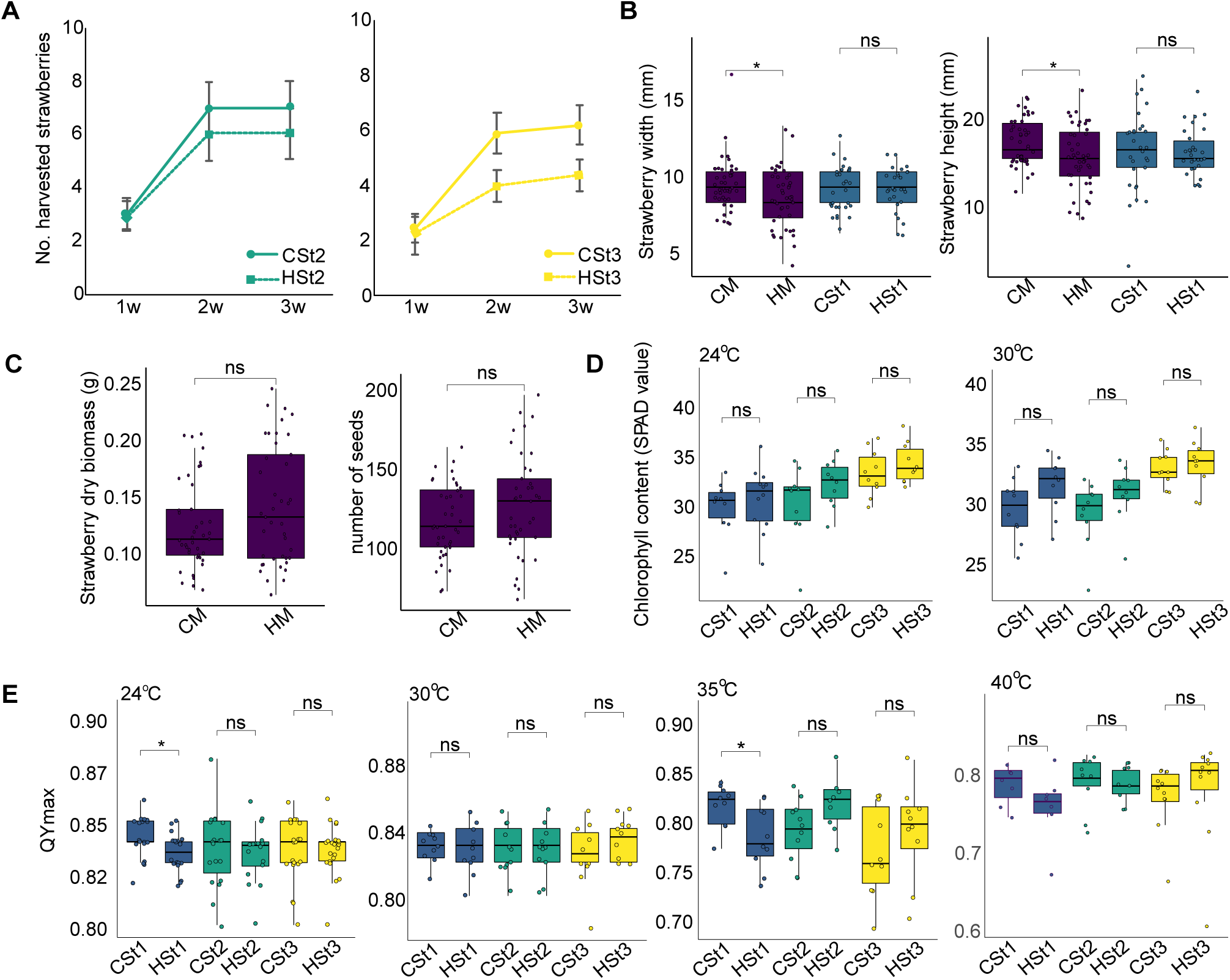
(A) Number of harvested strawberries during three consecutive weeks in M and St generations. (B) Strawberry size: width and height (mm). (C) Strawberry dry biomass and Number of seeds per fruit. (D) Total leaf chlorophyll content by the measurement of the Soil Pant Analysis Development (SPAD) values by chlorophyll meter of plant submitted to ascending temperature gradient (+5°C every 24h). (E) Maximum quantum yield of photosystem II (PSII) (QYmax) using chlorophyll fluorometer of plant submitted to ascending temperature gradient (+5°C every 24h All statistical analysis comparisons applied with Wilcoxon rank sum tests: *P ≤ 0.05; ns: no significant. HM: heat-stressed mother plant; CM: control mother plant; St1: first daughter plant from M; St2: daughter plant from St1; St3: daughter plant from St2.

**Figure S7.**
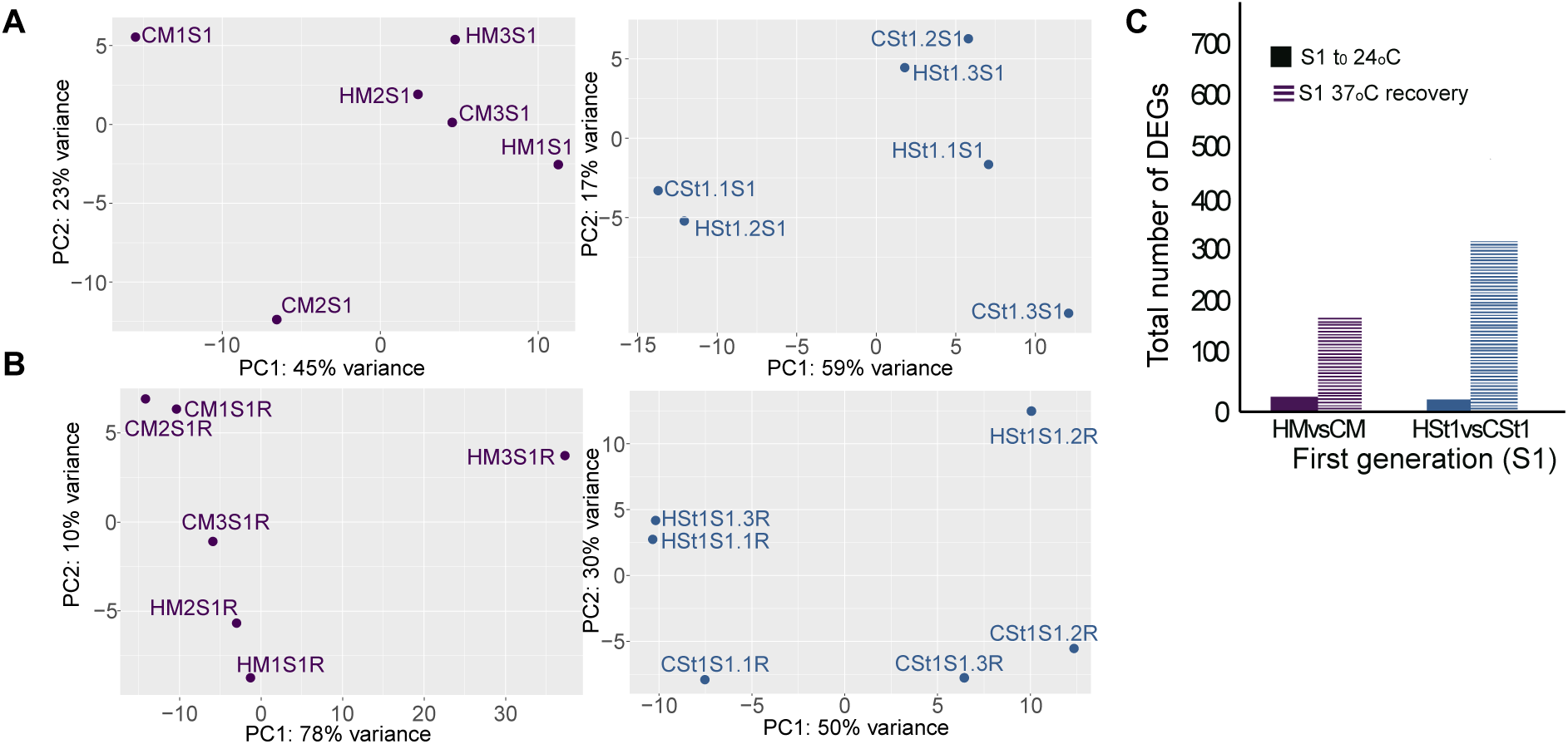
RNA-seq analysis to detect differentially expressed genes (DEGs) in the first-sexual generation (S1). PCA plots showing the variability among (A) control (B) and heat-stress (H) and at control conditions (24°C, t0); (b) Progeny after two days recovery (2d) from 24h 37°C treatment. (C) Total number of DEGs at t0 and 2d in the first progenies from HM, CM, St1 and St2. Plots and results were obtained by DESeq2 in the Galaxy platform.

## Supplementary tables in Excel file

**Table S1.** Bisulfite sequencing data quality analysis.

**Table S2.** RNA-seq analysis of differentially expressed genes (DEGs) after heat-stress in HM vs. CM.

**Table S3.** RNA-seq analysis of differentially expressed genes (DEGs) in the first daughter plants HSt1 vs. CSt1 in control conditions.

**Table S4.** RNA-seq analysis of differentially expressed genes (DEGs) in the daughter plants from the second clonal propagation level, HSt2 vs. CSt2, in control conditions.

**Table S5.** RNA-seq analysis of differentially expressed genes (DEGs) in the daughter plants from the third clonal propagation level, HSt3 vs. CSt3, in control conditions.

**Table S6.** DEGs related with a DMR in HM compared to CM.

**Table S7.** DEGs related with a DMR in HSt1 compared to CSt1.

**Table S8.** DEGs related with a DMR in HSt2 compared to CSt2.

**Table S9.** RNA-seq analysis of differentially expressed genes (DEGs) of first sexual generation from HMS1vsCMS1.

**Table S10.** RNA-seq analysis of differentially expressed genes (DEGs) of first sexual generation from HSt1S1vsCSt1S1.

**Table S11.** RNA-seq analysis of differentially expressed genes (DEGs) of first sexual generation from HM1S1vsCMS1 at 6h heat treatment.

**Table S12.** RNA-seq analysis of differentially expressed genes (DEGs) of first sexual generation from HMSt1S1vsCMSt1S1 at 6h heat treatment.

**Table S13.** RNA-seq analysis of differentially expressed genes (DEGs) of first sexual generation from HMS1vsCMS1 at 2d recovery from heat treatment.

**Table S14**. RNA-seq analysis of differentially expressed genes (DEGs) of first sexual generation from HSt1S1vsCSt1S1 at 2d recovery from heat treatment.

